# Ommochrome pathway knockout via CRISPR/Cas9 reveals sex-linked eye pigmentation and establishes a heritable genome-editing platform in Rhynchophorus ferrugineus

**DOI:** 10.64898/2026.07.26.740749

**Authors:** Hashel Bin Hraiz, Girlie Ann Agbayani, Ling Li, Jernej Jakše, Binu Antony, Khaled M.A. Amiri

## Abstract

The red palm weevil, *Rhynchophorus ferrugineus*, is the most economically destructive palm pest worldwide, threatening livelihoods, food security, and ecosystems across 49 countries. Weevil management currently relies predominantly on chemical insecticides, raising significant environmental and public health concerns. Despite its global agricultural importance, genetic approaches to pest management and the mechanistic basis of genome-editing strategies in *Rhynchophorus* remain largely unexplored. Here, we employed CRISPR/Cas9 genome editing to disrupt the *R. ferrugineus* ommochrome biosynthetic pathway — a multi-enzymatic metabolic cascade that converts tryptophan into ommochrome pigments, including brown, yellow, and red pigments. We targeted two key pathway components: the ATP-binding cassette transporter *white* and the heme peroxidase *cardinal*. Both genes were ubiquitously expressed, with peak expression levels in the gut, fat body, and head. Elevated transcript levels were observed across early, mid, and late pupal stages and in 0-, 1-, and 2-day-old adult males and females, consistent with the progression of eye pigmentation throughout the *R. ferrugineus* life cycle. Embryonic microinjection of a single guide RNA (sgRNA)–Cas9 ribonucleoprotein complex targeting *white* produced in the Generation-0 (G0) adults with a distinct, white-eyed phenotype with a brownish outer margin, in contrast to the black eyes of wild-type adults. Genome-edited *cardinal* mutant adults displayed a translucent, brownish-white-eyed phenotype, with white streaks that gradually transitioned to a persistent translucent reddish-brown eye coloration. Mutations in both genes were confirmed in G0 adults by genomic DNA sequencing. Mutant adults were crossed to generate heterozygous G1 (+/−), G2 (−/−, −/+, and +/+), and G3 lines (−/−) with genotypes verified as carrying 2-, 3-, 9-, and 13-nucleotide deletions. A stable, heritable eye-color phenotype was established in homozygous knockout (−/−) G3 lines for both *white* and *cardinal*, confirmed by unambiguous indel (insertions/deletions) genotyping. Inheritance analysis revealed that both genes are X-linked, following a classical Mendelian sex-linked pattern: paternal alleles are transmitted exclusively to daughters, while maternal alleles are inherited equally by both daughters and sons. This study establishes the first fully homozygous knockout strain in *R. ferrugineus* and, by characterizing sex-linked inheritance in a coleopteran system, advances our understanding of how CRISPR/Cas9 can be efficiently applied to destructive palm weevil species. The present study represents the first report of CRISPR/Cas9 genome editing in any weevil (Curculionidae), using *white* and *cardinal* as marker genes. These findings provide a valuable platform for functional genomics and genome engineering in *R. ferrugineus* and offer a translational framework for genome editing in the invasive South American palm weevil, *R. palmarum*, laying a solid foundation for the development of gene-drive strategies aimed at sustainable palm weevil population control.

## Introduction

Palm weevils (*Rhynchophorus* spp.) (Coleoptera: Curculionidae) are the most destructive invasive pests of economically important palm species worldwide, including coconut, oil, date, sago, and Canary Island palms. Among the nine *Rhynchophorus* species recognized as palm pests, the red palm weevil (RPW), *R. ferrugineus*, and the South American palm weevil (SAPW), *R. palmarum*, are the most damaging, collectively threatening coconut, date, oil, sago, and Canary Island palm cultivation across multiple continents (Hoddle et al., 2024; Hoddle et al., 2026). Both species are characterized by cryptic larval stages during which voracious feeding causes extensive internal damage to palm tissue, facilitating colonization by rot-inducing microorganisms and, if infestations go undetected, ultimately killing the host palm (Hoddle et al., 2026). Notably, *R. palmarum* is the sole confirmed vector of *Bursaphelenchus cocophilus*, the red ring nematode, which causes red ring disease — a lethal palm disease with no known cure (Hoddle et al., 2026). The RPW, also known as the Asian palm weevil, is classified as a transboundary quarantine category 1 pest by the United Nations Food and Agriculture Organization (FAO). Originally restricted to coconut and oil palm in Southeast Asia during the mid-19th century, *R. ferrugineus* expanded its invasive range dramatically in the late 20th century and now poses a severe threat to date palm, coconut, and Canary Island palm cultivation across the Middle East, North Africa, and the Mediterranean basin (Hoddle et al., 2024).

The RPW has attracted considerable global attention owing to its ongoing geographic expansion, now threatening palm ecosystems across 49 countries (Hoddle et al., 2024). Native to Southeast Asia and Melanesia, *R. ferrugineus* has been inadvertently introduced worldwide through the commercial trade of palm planting material over the past five decades. A recent FAO survey recorded RPW infestations on 53% of date palm farms in Saudi Arabia, with annual treatment costs estimated at USD 34.4 million. In recent decades, *R. ferrugineus* has undergone significant intracontinental range expansion, accompanied by rapid genetic adaptation to the extreme environmental conditions of the Middle East and Mediterranean regions. The invasion has substantially impacted oil palm and coconut production in Indonesia and Malaysia—the world’s two largest palm oil-producing nations—and has caused economic losses estimated at approximately €200 million in Europe alone in 2023, particularly in Italy, Spain, and France. In the Caribbean, RPW threatens coconut industries, has the potential to devastate national economies, and destroys royal palms, which are central to the region’s tourism landscape. Furthermore, RPW threatens palms of outstanding cultural and heritage significance, including those at UNESCO World Heritage Sites such as Socotra Island (Yemen) and iconic locations such as Cannes, southern France (Hoddle et al., 2024). In response to this escalating threat, the Consortium for Red Palm Weevil Control (C4RPWC) was established as a global programme to protect palms, livelihoods, and ecosystems worldwide (https://icarda.org/research/featured/protecting-palms-livelihoods-and-environment).

Current RPW management relies on costly integrated pest management (IPM) strategies that combine strict quarantine measures, early detection protocols, pheromone-based mass trapping, and targeted insecticide applications — all of which pose significant environmental, public health, and community risks. The indiscriminate use of insecticides in date palm plantations across the Middle East over the past two decades has resulted in pesticide residues in the environment and the emergence of insecticide-resistant weevil populations (Antony et al., 2019). While IPM has partially mitigated RPW impacts, the continued dependence on chemical control and the emergence of resistant populations underscore the urgent need for alternative, sustainable management strategies. Genetic control approaches offer an environmentally targeted and species-specific solution for managing insect pest populations (Arjunan and Thiruvengadam, 2026; Saini et al., 2026). Although functional genomics tools, including loss-of-function studies via RNA interference (RNAi), have been applied in RPW (Antony et al., 2019; Antony et al., 2021), field-level deployment of RNAi-based biopesticides remains constrained by the cryptic larval lifestyle and the practical difficulties of delivering dsRNA to infested palms (Sattar et al., 2024). The recent application of CRISPR/Cas9 in coleopteran pest management programmes has established the technology as a relatively efficient and cost-effective genetic tool (Li et al., 2025). Unlike RNAi, which mediates transient gene silencing, CRISPR/Cas9 introduces heritable, targeted mutations that can be stably transmitted across generations.

Despite the tremendous progress made in the last decade in insect genome editing, to date, only a handful of insect species have been targeted for CRISPR/Cas9-based genome editing, and this has been mainly focused on model insects, medically important insects, and key agricultural pests (Arjunan and Thiruvengadam, 2026; Saini et al., 2026). Notably, in Coleoptera, only four species have been attempted, namely the red flour beetle, *Tribolium castaneum* (Tenebrionidae) (Liu et al., 2025; Shirai and Daimon, 2020), Colorado potato beetle, *Leptinotarsa decemlineata* (Chrysomelidae) (Gui et al., 2020), Alligatorweed flea beetle, *Agasicles hygrophila* (Chrysomelidae) (Fu et al., 2025), and Western corn rootworm, *Diabrotica virgifera virgifera* (Chrysomelidae) (Chu et al., 2018). Importantly, the family Curculionidae, the most prominent family of beetles as agricultural pests and pollinators, with over 80,000 species reported to date (Bandeira et al., 2021), has not yet undergone CRISPR/Cas9 genome editing studies (Li et al., 2025). Despite the agricultural importance of *R. ferrugineus*, the genetic mechanisms underpinning its biology and the feasibility of genome-editing approaches in this species remain largely unexplored. Similarly, the application of CRISPR/Cas9 in *R. palmarum* has not advanced due to the lack of genomic, transcriptomic, and functional genomics resources (Hoddle et al., 2026). By contrast, a near-chromosomal-level genome assembly and multiple tissue transcriptomes for *R. ferrugineus* have recently been published, alongside functional validation studies, providing a resource base for the development of genetics-based control tools (Antony et al., 2024; Antony et al., 2016; Scieuzo et al., 2024; Sudalaimuthuasari et al., 2024). Nevertheless, a major bottleneck to genome editing in RPW is its cryptic life cycle, which complicates the collection of fertilized embryos and the determination of optimal timing and methods for embryonic microinjection. To date, no systematic methodology exists for weevil embryo microinjection, mutant population screening, or assessment of germline transmission efficiency — leaving critical gaps in the design of crossing schemes for stable line generation. A rigorous and systematic CRISPR/Cas9 editing framework for palm weevils would therefore provide the methodological foundation necessary for genetics-based biocontrol programmes.

To address these knowledge gaps, this study demonstrates, for the first time, CRISPR/Cas9-mediated genome editing in *R. ferrugineus*. We designed highly efficient single-guide RNAs (sgRNAs) with no predicted off-target effects and used high-fidelity (HiFi) Cas9 to target two key genes in the ommochrome biosynthetic pathway governing eye pigmentation. The first target, the ATP-binding cassette (ABC) transporter *white*, mediates intracellular transport of the intermediate precursor 3-hydroxykynurenine — synthesized from tryptophan — into pigment granules. The second target, the heme peroxidase *cardinal*, catalyzes the oxidative condensation of 3-hydroxykynurenine into ommochrome pigments. We first characterized the gene structure, tissue-specific and developmental expression profiles, and phylogenetic relationships of both *white* and *cardinal*. We then generated loss-of-function mutations in both genes via CRISPR/Cas9, characterized the resulting phenotypic and genotypic changes, and developed a systematic crossing strategy to establish stable homozygous knockout lines. Disruption of both genes markedly impaired the ommochrome biosynthetic pathway, resulting in distinct eye-color phenotypes. We further investigated the sex-linked inheritance of mutant eye color in *R. ferrugineus* through controlled mating experiments. Collectively, this study establishes a systematic methodology for embryo selection and microinjection, fills critical knowledge gaps in stable knockout line generation *via* crossing, and provides a foundation for the development of genetics-based control tools and future gene-drive programmes for palm weevil management.

## Materials and Methods

### Red palm weevil rearing

The red palm weevil (RPW) used in this study originated from a date palm field in Al Ain (UAE) in 2019 and was maintained in laboratory culture (25-27 °C; relative humidity ∼60 %; 14 h light and 10 h dark)—briefly, two male and two female adults were placed in a container containing molasses-soaked cotton wool. Cotton wool was laid out in layers to facilitate easier embryo collection. Containers were incubated at 25-27 °C for 3 to 4 days. Newly laid eggs were carefully collected, placed in a Petri dish, and checked daily to collect newly hatched larvae, which were added to distilled water as needed. After a week, the larvae were transferred to larger containers with synthetic medium and maintained on an artificial diet (Table S1) until they reached the late larval stage. The late larvae were transferred to processed sugarcane stems in the sixth week for pupation. RPW pupae were maintained on sugarcane stems, as described previously (Antony et al., 2021). The laboratory-reared RPW culture was considered a pure wild-type (WT) laboratory colony, free of admixture with other populations.

### RNA extraction and cloning of full-length *white* and *cardinal*

The gene structure and full-length *white* (Rferwhite) and *cardinal* (RferCar) sequences were retrieved from the RPW genome dataset at NCBI (acc nos. GCA_030347505.1), and oligonucleotide primers were designed (Table S2). Total RNA was extracted from whole-body tissue (∼30 mg) of laboratory-reared male and female RPW adults using the PureLink RNA Mini Kit (Thermo Fisher Scientific, USA), and genomic DNA contamination was removed with the PureLink DNase Set (Invitrogen) according to the manufacturer’s instructions. First-strand cDNA was synthesized using SuperScript IV Reverse Transcriptase (Thermo Fisher Scientific, USA) from 1 µg of total RNA. PCR was performed using the Applied Biosystems PCR System (Thermo) with the following conditions: 94°C for 5 min, followed by 35 cycles at 94°C for 15 sec, 60°C for 30 sec, and 72°C for 4 min, then a final extension at 72°C for 10 min. The PCR product was separated by agarose gel electrophoresis and purified using the Wizard® SV Gel and PCR Clean-up system (Promega). The purified DNA was ligated into the pGEM®-T easy vector and transformed into JM109 *E. coli* cells (Promega). Plasmid DNA was purified using the QIAprep Miniprep kit (Qiagen, Venlo, Netherlands), and the insert was sequenced in both directions (ABI 3500, Thermo) using primer-walking (Table S2). Sequences were aligned and annotated using BLASTx, and the results were further verified manually in Geneious *v*7.1.9 (Biomatters).

### Rferwhite and RferCar annotation, expression profiling, and gene structure analysis

Previously generated transcriptomes from different tissues of laboratory-reared and field-collected male and female RPW adults were retrieved (Antony et al., 2024; Scieuzo et al., 2024) and used for *white* and *cardinal* expression mapping. Transcriptome assembly and annotation were carried out as previously described (Antony et al., 2024; Gonzalez et al., 2021) using the Qiagen CLC Genomics Server (*v* 21.0.1). The cleaned RNA-seq reads were mapped to the *R. ferrugineus* genome (GenBank: GCA_030347505.2) using the ‘RNA-seq analysis procedure’ implemented in CLC. Further, the *white* and *cardinal* sequences were annotated and mapped to the *R. ferrugineus* genome (GenBank: GCA_030347505.2) by BLASTN search against the *R. ferrugineus* genome using Geneious *v*7.1.9 (Biomatters). The NCBI BLASTx homology search was used to verify the *white* and *cardinal* sequences and to check the open reading frames (ORFs). Gene expression levels were quantified and reported as reads per kilobase of transcript per million mapped reads (RPKM) and transcripts per kilobase of exon model per million mapped reads (TPM). The transcript levels were normalized to TPM values for *cardinal* and *white* using male vs. female *R. ferrugineus* transcriptomes. An expression heatmap was generated from transcript abundance values (TPM). Prior to visualization, TPM values were log₂-transformed as log₂ (TPM + 1). Expression values were averaged for each gene, body part, and sex across field and laboratory samples, excluding missing values. A heatmap was generated in Python using Seaborn (https://seaborn.pydata.org), with missing values masked and log-transformed expression values displayed in each cell.

The exon-intron positions of *white* and *cardinal* were mapped onto the *R. ferrugineus* genome (GenBank: GCA_030347505.2). The genomic regions were extracted, mapped, and manually aligned using MAFFT v7.38, which was also used for gene structure illustrations. The raw data of the *R. ferrugineus* tissue-specific transcriptome (Scieuzo et al., 2024) (Table S3) were assembled and cleaned. The reads corresponding to *white* and *cardinal* were visualized in the CLC Genomics Server using marked *white* and *cardinal* gene positions in the *R. ferrugineus* genome (GenBank GCA_030347505.2). To investigate the *white* and *cardinal* exon structures and genomic locations, the Qiagen CLC Genomics Workbench was used to extract genomic coordinates and exon annotations from the GFF file of newly annotated *R. ferrugineus white* and *cardinal* genes.

### Sequence alignment and phylogenetic analysis

Phylogenetic analyses were performed using *white* and *cardinal* amino acid sequences from *R. ferrugineus* and representative species from Coleoptera, Lepidoptera, Diptera, Orthoptera, and Hymenoptera. Multiple sequence alignments were performed using MAFFT v.7.45 with the auto (FFT-NS-1, FFT-NS-2, FFT-NS-i, or L-INS-i) option, depending on data size and strategy. The default parameters were applied, followed by manual trimming to remove gaps and ambiguous sequences. The auto algorithm and BLOSUM62 were used as the scoring matrix. Subsequently, Noisy *v*. 1.5.12.1 (Dress et al., 2008) (https://galaxy.pasteur.fr/) with default parameters was used on NGPhlyogeny. fr (Lemoine et al., 2019) to identify conserved regions of the alignment. The automatic model search was performed using ModelFinder 47, and the JTTDCMut+F+I+G4 and LG+I+G4 substitution models for *white* and *cardinal*, respectively, were determined as the best-fitting models according to the Bayesian information criterion (BIC). The resulting trimmed alignment contained 99 sequences with 2507 amino acid sites for the *white* and 93 sequences with 2745 amino acid sites for *cardinal*. Maximum-likelihood phylogenies were constructed with 1000 bootstraps in IQtree *v*1.6.12 (parameters: – bb 1000) (Nguyen et al., 2015), rooted in the Orthoptera lineage. The tree was visualized and edited with FigTree *v*1.4 (tree.bio.ed.ac.uk), colored, and finally edited with Adobe Illustrator (Adobe, CA, USA).

### Tissue-specific and relative expression analysis by RT-qPCR

Tissue-specific expression analysis was performed using cDNAs prepared from the antenna, gut, head, leg, thorax, and wings as mentioned earlier (Antony et al., 2018). First-strand cDNA was synthesized from ∼1.0 µg of total RNA using the QuantiTect reverse transcription kit (Qiagen, Germany), with genomic DNA removed using the gDNA wipeout buffer (Qiagen) as per the manufacturer’s instructions. RNA quantity and quality were assessed using a NanoDrop spectrophotometer (Thermo Fisher), and cDNA quality was verified by PCR amplification with the *tubulin* and *actin* genes (Antony et al., 2019) (Table S2). The PCR reactions were carried out using Q5 High-Fidelity 2X Master Mix (New England Biolabs, UK) with the following conditions: 95°C for 3 min; 35 cycles at 95°C for 30 s, 60°C for 30 s, and 72°C for 30 s, using a Veriti PCR System (Thermo). PCR products were evaluated on a 2.0% agarose gel containing HydraGreen™ Safe DNA Dye (ACTGene, NJ, USA) and visualized by electrophoresis with a 1kb DNA ladder (Thermo Fisher).

Expression levels of the *white* and *cardinal* genes were quantified by RT-qPCR across developmental stages (pupae and adults) and in different tissues, including the antennae, gut, fat body, head, legs, thorax, and wings. qRT-PCR was performed using FastSYBR® Green PCR Master Mix (Thermo Fisher) on the StepOnePlus Real-Time PCR System (Thermo Fisher). The thermal cycles program as followed was: 50°C for 20 s (pre-cycling), 95°C for 10 min (holding), followed by 40 cycles of 95°C for 15 s, 60°C for 35 s, and a melt curve analysis was performed with 95°C for 15 s, 60°C for 1 min, 95°C for 30 s, and 60°C for 15 s. The reactions were performed using three biological and three technical replicates. Relative gene expression in the antennae was compared with that in other tissues. Data were calculated using the comparative 2-ΔΔCт method (Livak and Schmittgen, 2001). The RPW *tubulin* and *β-actin* (Table S2) were used as endogenous controls (Antony et al., 2019).

### CRISPR/Cas9-mediated editing of *white* and *cardinal*

#### Synthesis of single-guide RNA

The principle of 5′-N20<u>NGG</u>-3’ (protospacer-adjacent motif (PAM) underlined) was followed to design the sgRNA recognition site. Dual sgRNAs were designed to target exons 2 and 6 of *white* and exons 3 and 5 of *cardinal*, respectively. The potential off-target effects were analyzed using Chopchop (https://chopchop.cbu.uib.no/) and CRISPOR (https://crispor.gi.ucsc.edu/crispor.py), followed by manual verification. The CRISPR-Cas9 guide RNA design checker provided by Integrated DNA Technologies (IDT) (Leuven, Belgium) was used to assess the on-target potential of protospacer designs. Those gRNAs with the highest on-target scores, indicating the best predicted editing performance at the intended target site, were selected and synthesized using Alt-R gRNA modifications (IDT). The Alt-R™ *Streptococcus pyogenes* (S.p.) Cas9 High Fidelity (HiFi) Cas9 Nuclease V3 (500 μg) was purchased from IDT.

### *R. ferrugineus* egg collection, embryo microinjection

A customized protocol was developed for fertilized egg collection, embryo microinjection, and subsequent weevil rearing. For adult mating and subsequent egg collection, a black, aerated plastic box with three compartments was used. For egg collection, seven males and ten females were placed in each compartment of an oviposition box with a layer of freshly prepared semi-solid molasses agar. The RPW adults were kept undisturbed overnight (25–27 °C) to allow mating. The following morning, the bottom plate was replaced with a fresh layer of molasses agar, and the mated females were allowed to oviposit for 2-4 hours. The newly laid eggs were carefully aligned on a glass slide, and microinjection was performed into the posterior or germ band region of embryos using a dual sgRNA injection strategy. The injection mix contained 300 ng/uL of each sgRNA and 300 ng/uL of HiFi S.p. Cas9 (IDT). The mixture was incubated at 37 ^°^C for 30 min to allow formation of the ribonucleoprotein (RNP) complex, and microinjection was completed within 30 min of egg collection. Embryo microinjection was performed using a semi-automated Sutter microinjection system (Novato, CA, USA) mounted on an Olympus SZX10 stereomicroscope (Tokyo, Japan) with a borosilicate glass microinjection capillary needle (100 mm length; 1.0 mm outer diameter and 0.58 mm inner diameter with filament polished). An Eppendorf Microloader (Eppendorf, Germany) was used to load the injection fluid into the needle. After injection, the embryos were transferred onto a wet filter paper placed in a Petri dish and incubated at 25–27 °C in a growth chamber until hatching. Upon hatching, the larvae were transferred to a plastic box containing an artificial diet (Table S1) and followed the rearing protocol described above. RPW pupae were maintained on sugarcane stems, as described previously (Antony et al., 2016).

#### Mating experiment and raising a homozygous line

The newly emerged adult G0 RPW males and females were kept separately, without mating, until their genotypes were determined by PCR amplification and sequencing of the genomic regions flanking each sgRNA target site using specific primers (Table S2). Genomic DNA was extracted from the mid-leg (dissected tibia and tarsi) of each G0 adult using the Quick-Tissue/Insect genomic Microprep Kit (ZymoResearch, MA, USA). PCR amplification [95 °C for 5 min, 35 cycles of 95 °C for 1 min, 60 °C for 30 s, and 72 °C for 30 s; and one cycle at 72 °C for 10 min] was carried out using the Q5 High-Fidelity 2X Master Mix (New England Biolabs). The PCR products were evaluated as previously described, purified using the Wizard® SV Gel and PCR Clean-up system (Promega), and sequenced on an ABI 3500 genetic analyzer (Thermo). G0 individuals with a distinct phenotype (eye color variant) and sequencing-verified genotype were selected and crossed with WT in a one-to-many mating scheme (G01:WT5). Progeny from successfully mutated G0 individuals were retained as the G1 generation. Within the G1 generation, individuals with distinct phenotypes and confirmed genotypes were selected for single-pair sibling crosses. The resulting G2 offspring were segregated and screened for homozygous mutants (−/−) by DNA sequencing, and the remaining offspring were retained for future line stabilization. A single G2 sibling pair with the same indel (insertions/deletions), verified by DNA sequencing, was selected for inbreeding to establish a stable homozygous mutant line (−/−) in the G3 generation.

### *White* and *cardinal* expression analysis in the mutant line

Homozygous *white* (3-nt deletion) and *cardinal* (13-nt deletion) mutant RPW adults were selected for expression analysis and compared with WT adults. Gut and head tissues were dissected from 2-5 days old adults, and RNA extraction and cDNA synthesis were performed as described previously. PCR amplification of the gRNA target regions in WT and mutant *white* and *cardinal* strains was performed using specific primers (Table S2), and the products were sequenced in both directions. qRT-PCR was performed using FastSYBR® Green PCR Master Mix (Thermo) on the StepOnePlus Real-Time PCR System (Thermo), following the aforementioned method, with *tubulin* and *actin* as reference genes (Antony et al., 2019), and WT was used as the calibrator.

### Mating frequency and sex-linked inheritance of *white* and *cardinal* mutants

We tested whether the *white* and *cardinal* indel knockout homozygous lines affected mating and fecundity by conducting a series of mating experiments with unmated homozygous males and WT virgin females. Homozygous mutant (−/−) *white* and *cardinal* adult males were crossed with WT females at a 1:5 male-to-female ratio for 3 weeks. Eggs were collected twice a week and reared to adulthood. The control mating experiment was performed with WT male and female individuals under the same laboratory conditions. Eggs were collected from each cross twice weekly and reared to adulthood. Genomic DNA was extracted from the mid legs of surviving adults using the Quick-Tissue/Insect Genomic Microprep Kit (ZymoResearch). The genotype was determined by PCR amplification and sequencing of the flanking regions of each gRNA target site using specific primers (Table S2).

To analyze the inheritance pattern of *white* and *cardinal* alleles in *R. ferrugineus,* homozygous (verified through Sanger sequencing) *white* or *cardinal* mutant females (w−/w− or c−/c−) or males (w−/Y or c−/Y) were crossed with virgin WT females (W^+^/W^+^ or C^+^/C^+^) or males (W^+^/Y or C^+^/Y). The eye color of each newly emerged male and female progeny (0-1 day old) was recorded, and the resulting progeny were genotyped. Some of the sequences obtained showed ‘mixed base calls’ across the entire region (edited and unedited alleles) in our CRISPR-edited RPW samples. These genotypes were further verified through TA cloning using the Zero Blunt TOPO PCR cloning kit (Invitrogen). The gel-purified PCR product was ligated into the pCR™4Blunt-TOPO™ vector, transformed into DH5-α *E. coli* competent cells, and individual bacterial colonies were picked (each containing only one specific allele). The isolated plasmids (QIAprep Spin Miniprep Kit, Qiagen) were sequenced individually to obtain clean traces of each allele.

To further confirm sex-linked inheritance of *cardinal* in *R. ferrugineus*, a virgin heterozygous female carrying the 13-nt deletion (C^+^/c^−^) (verified through Sanger sequencing) was crossed with a WT male (C^+^/Y). Thereafter, the eye color of each newly emerged progeny (0-1 day old) was recorded and genotyped using specific primers (Table S2). Furthermore, the ORF of *cardinal* was amplified and sequenced from a hemizygous *cardinal* mutant (−/−) with the 13-nt deletion. We performed the cryptic splice site prediction on the *cardinal* sequence at the 13-nt deletion site using MaxEntScan (http://hollywood.mit.edu/burgelab/maxent/Xmaxentscan_scoreseq.html) to predict a new cryptic GT donor site created by the 13-nt deletion (Yeo and Burge, 2003).

#### Data analysis

For the qRT-PCR experiments, relative expression levels were calculated as fold changes, and the mean fold changes (2^-ΔΔCT) were calculated in Excel (Microsoft Corporation, USA). CT values were collected from three biological replicates, each with three technical replicates. One-way analysis of variance (ANOVA) was used to assess significant differences among the experimental groups for qRT-PCR, followed by multiple-comparison tests with the least significant difference (LSD) test (*p* < 0.05) using SPSS *v*24 (IBM SPSS Statistics, NY, USA).

## Results

### RferWhite and RferCardinal cloning, sequence analysis, and phylogeny

The open reading frame (ORF) of the *white* transcript was identified from a 2,040-bp cDNA sequence retrieved from RPW gut and head transcriptomes and confirmed by PCR amplification (Figure S1). The ORF encodes a protein of 679 amino acids (aa) with a theoretical isoelectric point (pI) of 5.84 and a molecular weight (Mw) of 76,338.12 Da. Based on BLASTp (NCBI) searches, *R. ferrugineus white* (RferWhite) shares 80.70% amino acid sequence identity with the rice weevil, *Sitophilus oryzae white* (GenBank: XP_030764169.1), which has the highest identity in the NCBI non-redundant (nr) protein sequence database. The conserved protein domain analysis identified RferWhite as a member of the Eye Pigment Precursor Transporter (TIGR00955) family protein (transport and binding proteins) (EC 7.6.2.6). Searches against the NCBI conserved protein domain (accession, cl36780) (Wang et al., 2023) and the KEGG Orthology (KO) database revealed molecular functions represented in terms of functional orthologs (K21396). *White* gene model annotations retrieved from the FlyBase consortium (https://flybase.org/reports/FBpp0070468) for *Drosophila melanogaste*r (∼55.35 % identity) (NCBI: NP_476787.1) were used to predict the conserved motifs and protein domains of RferWhite, which identified the following domains: pigment precursor permease (IPR005284), ATP-binding cassette, ABC transporter-type domain profile (IPR003439), P-loop containing nucleoside triphosphate hydrolase (IPR027417), AAA+ ATPase domain (IPR003593), ABC transporters family signature (IPR017871), and ABC-2 type transporter (IPR043926 and IPR013525) (Figure S1a).

The *R. ferrugineus cardinal* (RferCar) transcript contains a 2,400 bp ORF (Figure S2), encoding a protein of 799 aa with a pI/Mw: 8.39/ 91541.99 Da, respectively. RferCar shares 67.85% identity with the rice weevil, *S. oryzae cardinal* (GenBank accession XP_030755866.1). The conserved protein domain analysis identified RferCar as a member of the animal haem peroxidase family (pfam03098) (EC: 1.11.1.-), which was further supported by a search against NCBI conserved protein domains (accession, cl38107) (Wang et al., 2023). Gene model annotations retrieved from the FlyBase consortium for *D. melanogaste*r (∼44.00% identity) (NCBI: NP_651081.1) were used to predict RferCar motifs and identified conserved protein domains, including haem peroxidase domain superfamily (animal type) (IPR037120), animal haem peroxidase superfamily profile (IPR019791), and haem-dependent peroxidases (IPR010255) (Figure S2a).

The *white* and *cardinal* sequences obtained from publicly available transcriptomic and genomic projects for various insect orders were used to construct the phylogenetic tree. Both RferWhite and RferCar were grouped within a major coleopteran lineage and clustered in the Curculionid clade (Figure 1). RferWhite was closely related to the rice weevil, *S. oryzae* ABC transporter G family member 36-like (GenBank XP_030764169.1) (Figure 1). Similarly, RferCar was closely related to the rice weevil, *S. oryzae* haem peroxidase (GenBank XP_030755866) (Figure 1).

**Figure 1.**
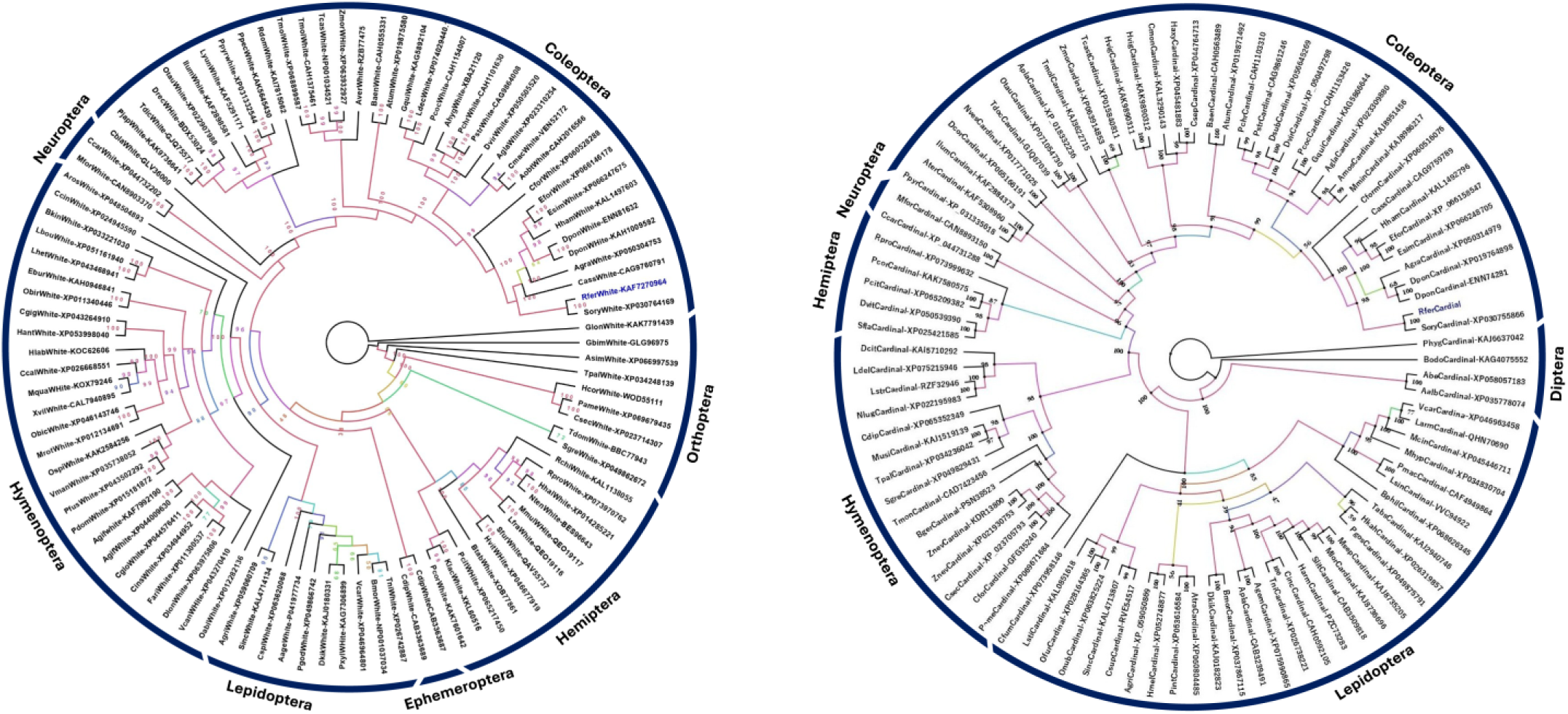
Maximum-likelihood consensus tree of *White* (left) and *Cardinal* (right) from insects. The tree was built from the alignment of amino acid sequences of *white* and *cardinal* representatives from neuropterans, coleopterans, orthopterans, hymenopterans, hemipterans, dipterans, and ephemeroptera. The orthopteran clade was used as an outgroup. The major insect orders are indicated with blue arcs. Numbers on the branches are bootstrap values (1000 ultrafast bootstrap (UFBoot) replicates). The phylogenetic tree was visualized using FigTree (http://tree.bio.ed.ac.uk/software/figtree/), and branch colors were based on bootstrap values. Scale = 3.0 amino acid substitutions per site. The sequences were retrieved from the GenBank database, and their accession numbers are shown alongside the taxon names.

### RferWhite and RferCar annotation, expression mapping, and gene structure analysis

We mapped *RferWhite* and *RferCar* expression in *R. ferrugineus* adult male and female tissue transcriptomes (gut, fat body, head, thorax, antennae, wings, and legs) (Table S3a-h). Both genes were then evaluated based on their expression levels, measured as transcripts per million (TPM) in the male and female tissue transcriptomes. The results indicated that RferWhite and RferCar expression patterns were similar between males and females (Figure 2A). Both *white* and *cardinal* were highly expressed in the gut and fat body of both sexes, followed by the head and wings (Figure 2A). The exon-intron structures of *white* and *cardinal* were retrieved from the RPW genome assembly (Sudalaimuthuasari et al., 2024) and mapped onto RPW tissue transcriptomes. By mapping RNA-seq reads to the *R. ferrugineus* genome, the tissue-specific expression profiles of the *white* and *cardinal* in *R. ferrugineus* were established. Analysis of total exon read counts for RferWhite and RferCar in both male and female tissues further confirmed that both genes showed the highest expression in the gut, fat body, head, and wings (Figure 2A).

**Figure 2.**
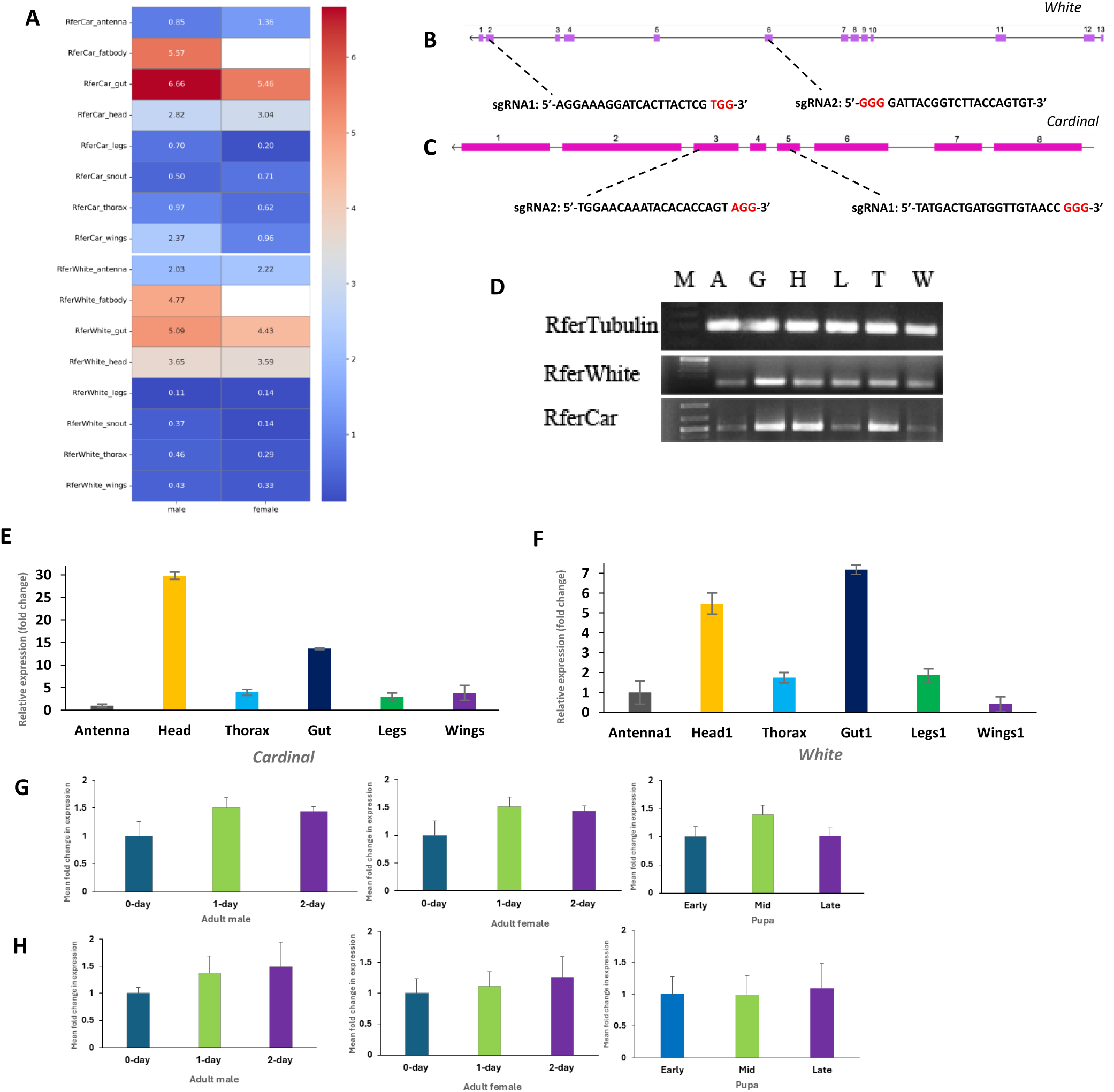
A. Expression profiles of RferWhite and RferCar in the male and female tissue transcriptomes. Expression levels of RferWhite and RferCar in the RPW tissue transcriptomes (Table S2) are represented as heat maps based on the log2-transformed transcripts per million (TPM) values. **B & C.** Schematic representations of the RferWhite (*white*) and RferCar (*cardinal*) genes in *R. ferrugineus*. The boxes represent the exons (numbered) encoding the protein, and the intron sequences are represented in dashed lines. The genomic region in the exon corresponding to the guide RNA target sites (black) and PAM sites (red) is highlighted. **D.** Tissue-specific expression analysis of RferWhite and RferCar. Tissues used are indicated as A (antennae), G (gut and fat body), H (head), L (leg), T (thorax), W (wings), and M (marker). Primer details are provided in Table S2. **E & F**. The original gel image (with DNA ladder) is provided in Figure S3. Tissue-specific relative expression levels of the *white* of *cardinal* gene in *R. ferrugineus*. Relative expression levels (mean ± SEM, n = 3) were calculated using the ΔΔCt method and normalized to the geometric mean of the *tubulin* and *actin* reference genes. Antennae were used as the calibrator tissue (1.0×). **G**. Relative expression levels of RferWhite in RPW pupae, male, and female adults (fold changes compared to the expression level on day 0). **H**. Relative expression levels of RferCar in RPW pupa, male, and female adults (fold changes compared to the expression level on day 0).

The structural positions of *white* and *cardinal* on the X-chromosome were characterized, and their organization was analyzed. *R. ferrugineus* possesses a diploid chromosome number of 2n=22, featuring a sex-determination system with one acrocentric X chromosome and one dot-shaped Y chromosome (Xyp parachute sex-determination system) (Al-Qahtani et al., 2014; Sudalaimuthuasari et al., 2024). Using the near-chromosomal level genome of RPW (Sudalaimuthuasari et al., 2024), and Chromosome =“ NW_027472586” (rfm7-Chr7; 9402771..9419283), we identified the *white* gene (LOC143200194), which spans 16,513 nt with 18 exons and 11 introns, and the aligned mRNA sequence length of 4,049 nt. The *white* coding sequence (CDS) spans 3,963 nt and encodes a deduced amino acid (aa) sequence of 1,320 aa. However, the *white* ORF retrieved from the gut tissue transcriptome (NCBI SRR27695095) was 679 aa, and this discrepancy in the predicted ORF length was further confirmed by PCR amplification and sequencing (Figure S1).

To further investigate the discrepancy, the predicted 1,320-aa fragment was analyzed using NCBI SmartBlast against the UniprotKB/Swiss-Prot database and mapped to the *D. melanogaster* genome in FlyBase. We found that the reported 1-648 aa sequence corresponds to a partial sequence of the *scarlet* protein, encoded by an autosomal gene on the third chromosome (73A3-4). *Scarlet* is a subunit of another ATP-binding cassette (ABC) transporter (Shamim et al., 2014). The *white* dimerizes with *scarlet* to transport pigments into pigment granules. Nevertheless, *D. melanogaster white* encodes a 687-aa protein (NCBI: NP_476787) and is positioned on ChrX (NC_004354.4) (GCF_000001215.4). Due to errors in the RPW genome assembly and annotation (Sudalaimuthuasari et al., 2024), the *white* gene LOC143200194 (positions 9,402,769-9,419,283) has been incorrectly assigned to the autosome (rfm7) (Sudalaimuthuasari et al., 2024). Therefore, we reassembled and reannotated the RPW genome and generated a revised gene model for RferWhite in the recently published RPW genome (https://bipaa.genouest.org/sp/rhynchophorus_ferrugineus/) (NCBI BioProject database (BioProject: PRJNA967181, Biosample: SAMN34578471, Accession: JASGYX000000000). We confirmed the correct RferWhite mRNA sequence length of 2302 nt, the ORF-spanning length of 2,040 nt, and the deduced amino acid sequence of 679-aa. We identified a *white* gene of 13,965 nt (coordinates 44560..58524) on scaffold 240, with 13 exons and 13 introns (Figure 2B). The reannotated *R. ferrugineus* genome assembly rfMv2 (Sudalaimuthuasari et al., 2024) available at the BIPPA database mapped RferWhite (gene ID: g26909) in the genome coordinate JASCQM010000007.1 9406736..9419227 (12,492 bp) with an ORF-spanning length of 2,040 bp (Figure S1), with 13 exons and 13 introns (Figure 2B).

Using the NCBI genome of RPW (GCA_030347505.2) and the X-chromosome = “NW_027472584.1” (rfm5 – Chr5) complement (19465423..19468630) (NCBI JASCQM010000005.1), we identified a *cardinal* gene length of 3,208 nt with 8 exons and 9 introns (Figure 2C). The predicted mRNA sequence length was 2,677 nt (XM_076406331.1), and the ORF length was 2,400 nt, with the deduced amino acid sequence of 799-aa (Figure S2).

### Tissue-specific and relative expression analysis by RT-qPCR

The RferWhite and RferCar were ubiquitously expressed across all examined tissues as confirmed by RT-PCR (Figure 2D). Based on RT-qPCR analysis, RferWhite and RferCar showed the highest expression in head and gut tissues (*p* < 0.001) compared to the other tissues (Figure 2E and 2F). We further quantified the relative expression of both *white* and *cardinal* in pupal and adult RPW using RT-qPCR. While both *white* and *cardinal* showed a progressive increase in expression levels from 0-day to 2-day-old adults, their expression increased differently during pupal stages (Figure 2G and H).

### CRISPR-Cas9-mediated mutagenesis *of White* and *cardinal*

We aimed to disrupt the ommochrome pathway (Figure 3A) using CRISPR/Cas9 genome editing in *white* and *cardinal* in RPWs. Two sgRNAs were designed to target exon 2 and 6 of *white* (Figure 2B). A mixture of both sgRNAs (300 ng/uL each) with a high-fidelity (HiFi) *Streptococcus pyogenes* (S.p.) Cas9 nuclease (300 ng/uL) was assembled to form a ribonucleoprotein (RNP) complex, which was injected into the freshly laid RPW eggs (G0). In total, 447 eggs were injected with the RNP complex, and the injected G0 larvae, pupae, and adults were maintained for phenotypic analysis. Of the 447 injected eggs, 48.54% hatched into first-instar larvae and only 22 individuals reached adulthood (Figure 3B), while the remaining individuals died during development. Among the 22 adult G0 RPWs, 17 (77.27%) were males, and five (22.72%) were females. Among G0 adults, we observed a distinct white-eye phenotype in one (5.88%) male and one (20.00%) female, which was further confirmed by Sanger sequencing. Overall, mutant males and females accounted for 0.20% and 0.21% of the 447 injected embryos, respectively (Figure 3B). The *white* mutant RPW adults exhibited a clear, translucent white eye with a brownish, outer-margined phenotype, whereas WT adults showed black compound eyes (Figure 3C and D). During development, the mutant eye color changed to brownish-white and eventually developed a persistent translucent reddish-brown eye-color phenotype (Figure 3D). In addition, the whole-body coloration changed from reddish-brown in WT to pale yellow in *white* mutant adults (Figure 3C).

**Figure 3.**
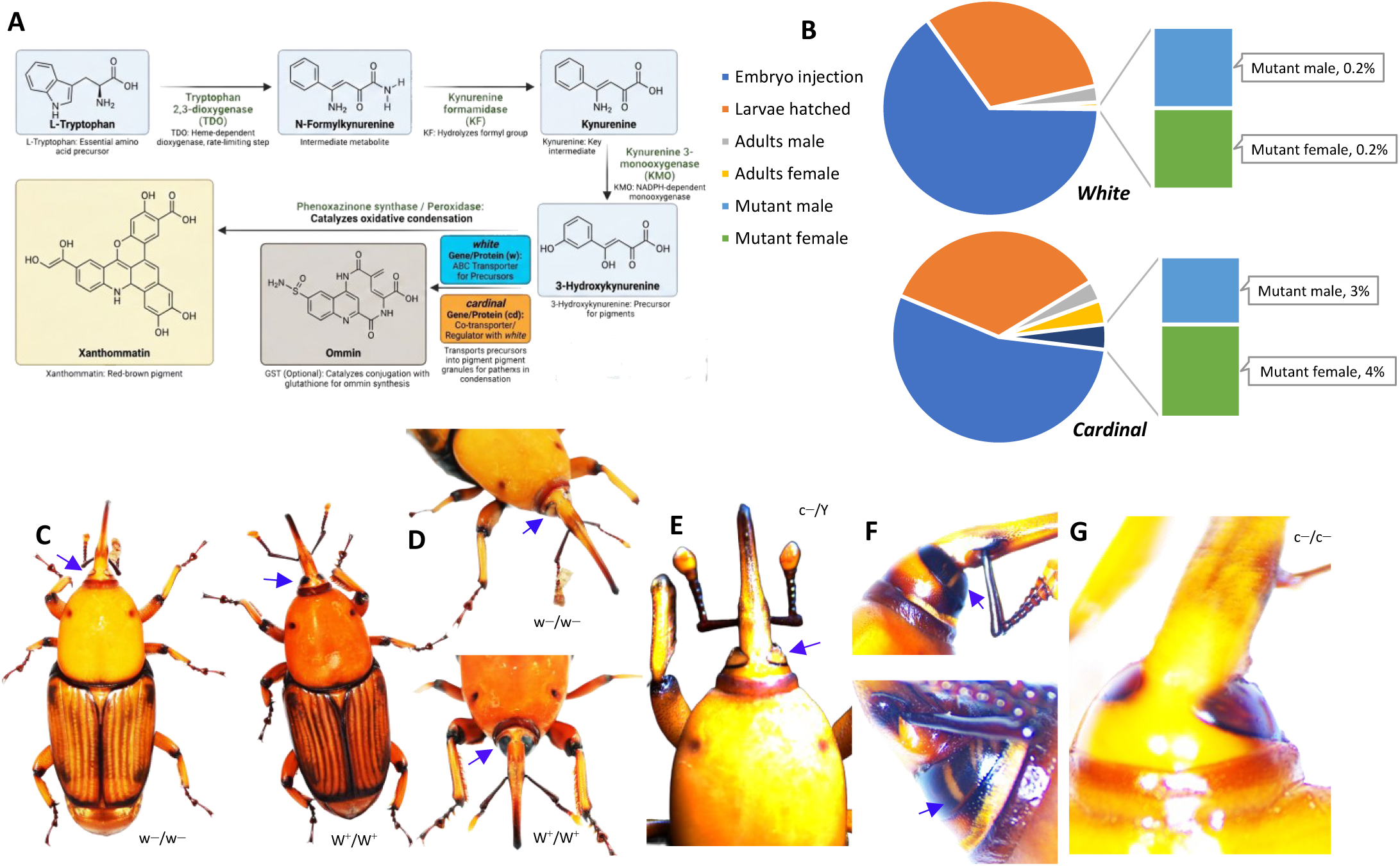
Different eye pigmentation phenotypes developed in the RferWhite and RferCar mutant adults. **A**. Schematic representation of the ommochrome pathway in *R. ferrugineus*. Substrates and enzymes are indicated in blue-grey and green, respectively—pathway direction indicated by an arrow. The *white* (w) and *cardinal* (cd) alleles were selected for genome editing and assessed for functional role, as indicated in the figure. **B**. A pie chart represents outcomes and mutant yield following embryo microinjection. Slices represent the proportions of injected embryos, successfully hatched larvae, emerged adults, and identified mutant males and females. **C**. Eye pigmentation and body color in wild-type (W^+^/W^+^) and *white* mutant (w−/w−) *R. ferrugineus* adults (0-day). The mutant exhibited white eyes (arrow), whereas WT adults had clear, black compound eyes. **D**. Eye pigmentation changes to translucent white color (arrow) in the *white* mutant (w−/w−), compared to WT adults (W^+^/W^+^). **E**. Eye pigmentation of *cardinal* mutant (c−/Y) *R. ferrugineus* adult (0-day). **F**. *Cardinal* mutant (c−/Y) exhibiting a black compound eye with a white streak (arrow) on the left eye, and a white streak on the right eye (arrow). The adults were photographed at the 0-day and 2-day adult stages. **G**. Eye pigmentation changes to translucent reddish-brown eye color in *cardinal* mutant adult (5-day-old).

Two RferCar sgRNAs were designed targeting exon 3 and 5 (Figure 2C). Both sgRNAs (300 ng/uL each) were mixed with Cas9 (300 ng/uL) to form an RNP complex and injected into freshly collected RPW eggs. In total, 370 eggs were injected with the RNP complex, and the injected G0 larvae, pupae, and adults were maintained for phenotypic analysis. Of the 370 injected eggs, 64.32% of the first-instar larvae hatched; only 46 RPWs reached adulthood (Figure 3B), and the rest died. Among the 46 G0 adults, 21 (45.65%) were males, and 25 (54.34%) were females. Among G0 adults, we observed a distinct *cardinal* phenotype in 11 (52.38%) males and 15 (60.00%) females, which was further confirmed by Sanger sequencing. Overall, mutant males and females accounted for 3.0% and 4.0% of the 370 injected embryos, respectively (Figure 3B). The *cardinal* mutant adults (G0) exhibited a translucent, brownish-white eye (Figure 3E) and a white-streaked eye (Figure 3F), which gradually developed into a translucent reddish-brown eye (Figure 3G). In contrast, WT adults displayed black compound eyes.

The *white* and *cardinal* mutations were further validated by PCR amplification and sequencing of the genomic DNA (gDNA) flanking the sgRNA1 and sgRNA2 target sites (see Table S2 for primer sequences). The presence of mutations was confirmed by mixed sequence peaks (mixed base calls) upstream of the protospacer adjacent motif (PAM) site on the sense strand due to non-homologous end joining (NHEJ) repair errors, which often result in indels (Figure 4A and B). Deep analysis of these positive mutants using the Inference of CRISPR Edits (ICE) platform (https://ice.editco.bio/) identified indels at the target sites (Figure 4B). ICE analysis identified the cut-site region, indicated by the vertical dotted line in the chromatogram (Figure 4B), when compared to the WT control samples. Hereafter, we followed a distinct crossing scheme to establish homozygous lines, as outlined in Figure 4A. Briefly, positive G0 mutants were outcrossed to WT individuals in a one-to-many mating scheme (1:5 ratio). Around 100 eggs were collected, and larvae were reared to adulthood (G1). Some newly emerged G1 adults showed apparent white eyes with a brownish outer margin, suggesting the germline transmission of the white mutations. Further, sequence analysis of gDNA flanking the sgRNA target sites and multiple sequence alignment with the WT sequence confirmed the CRISPR-induced indels (Figure 4B and 4C). The multiple sequence alignment inferred an indel present in the edited G1 mutant (Figure 4B). The G1-mutant adults carrying the same indel were selected for a single-pair sibling cross, and the collected eggs (G2) were reared to adulthood (Figure 4A). We obtained the expected Mendelian segregation ratio of 1:2:1 (+/+: +/−: −/−) with 25 % homozygous mutants carrying 3-nt and 9-nt deletions in exon 2 of RferWhite (sgRNA1 site), and a 2-nt deletion in exon 6 (sgRNA-2 site), resulting in a distinct *white* eye phenotype. Further, each homozygous line was used for a single-pair sibling cross (3-nt x 3-nt and 9-nt x 9-nt), and the collected eggs (G3-line) were all homozygous (−/−) and were maintained for further analysis. The RferWhite homozygous line (9-nt indel) chromatogram traces clearly depict the indel, followed by continuous base calls (Figure 4C).

**Figure 4.**
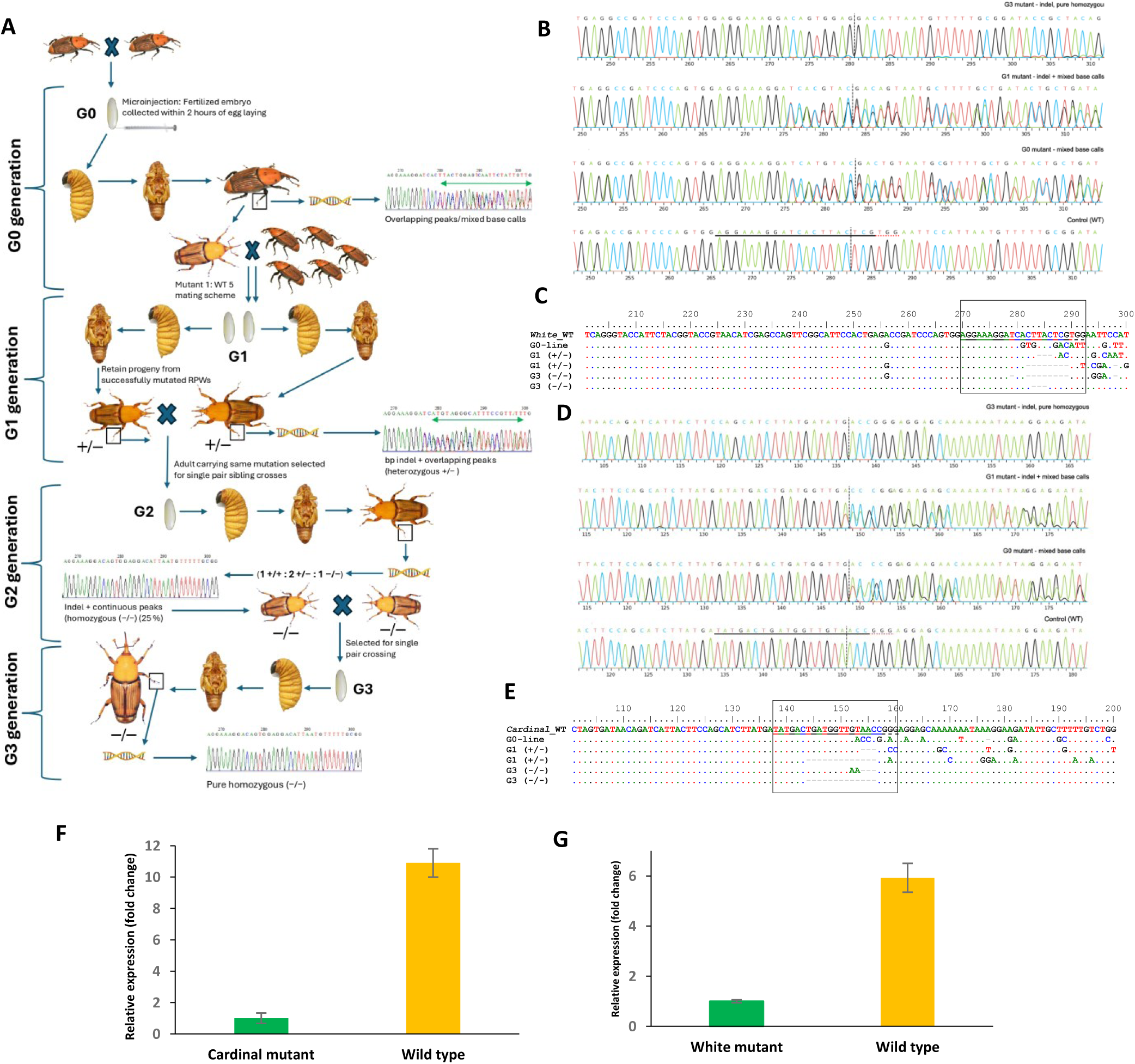
Schematic representation of embryo collection, sgRNA/Cas9 RNP complex microinjection, germline-derived G0-line, and the breeding scheme used to establish G1, G2, and G3 homozygous mutant lines of *R. ferrugineus*. **B & D**. Sanger sequence chromatogram alignment of RferWhite (B) and RferCar (D) mutants (heterozygous G0 and G1, and homozygous G3) compared to the wild-type (WT). This alignment of base calls from both the control and mutant samples in the AB1 files was visualized in ICE *v*4 https://ice.editco.bio/. The horizontal, black-underlined region represents the guide sequence, and the red-dotted underline indicates the PAM site. The vertical black dotted line represents the actual cut site. **C** & **E**. Multiple sequence alignment of RferWhite (C) and RferCar (E) regions corresponding to the guide RNA target sites (underlined) and the PAM site (dotted line). For the RferWhite G0 mutant with mixed base calls at the gRNA region, 3-nt and 9-nt indels identified in G1, G2, and G3 mutants are shown. For the RferCar G0 mutant with mixed base calls at the gRNA target region, 3-nt and 13-nt indels in the G1, G2, and G3 mutants are shown. The gRNA target region is highlighted in a square box. The number of deleted nucleotide sequences is shown as a grey dashed line in the multiple sequence alignment. **F & G**. Comparison of RferCar (left) and RferWhite (right) expression levels in gut tissue of mutant and WT RPWs. The mean fold change in gene expression relative to multiple endogenous controls (*tubulin* and *β-actin*) is shown, with SEM as error bars. A significant difference in RferCar expression was observed between mutant and WT, as based on one-way ANOVA (*p* = 0.001).

The *cardinal* mutations were verified by DNA sequencing of the flanking regions of sgRNA1 and sgRNA2, which show mixed peaks (mixed base calls) upstream of the PAM site (sense strand), indicating the presence of mutations that often result in indels (Figure 4D). Deep analysis of these positive mutants using ICE analysis identified an indel at the target site (Figure 4E). The ICE analysis predicted the region of the Cas9 cut site (Figure 4D) relative to WT control samples. We followed the aforesaid crossing scheme to raise a homozygous line (Figure 4A). The newly emerged G1 adults showed brownish-white-eyed, translucent brown-eyed, and white-streaked-eyed phenotypes, suggesting that the *cardinal* mutation in G0 could be inherited in the germline. Further, sequence analysis of the gDNA flanking sgRNA1 and sgRNA2 target sites, which yielded mixed base calls, and a multiple sequence alignment with the WT confirmed the CRISPR-induced indels (Figure 4E). The G1 mutant adults carrying the same indel were selected for single-pair sibling crosses, and the resulting eggs (G2) were reared to adulthood (Figure 4A). We obtained the expected 1:2:1 (+/+: +/−: −/−) ratio. A homozygous (−/−) line carrying 3nt and 13nt deletions in exon 5 of RferCar (sgRNA1 target site) (Figure S2) exhibited a distinct white-eye and white-streak eye phenotype, which developed into persistent translucent-brown. Further, each homozygous line (3-nt x 3-nt and 13-nt x 13-nt) was chosen for single-pair sibling crosses, and all G3 adults were confirmed as homozygous mutants and maintained for further analysis (Figure 4A).

### *White* and *cardinal* expression analysis in the mutant line

For the RferCar homozygous (−/−) G3 mutant carrying a 13-nt indel, the relative expression level (fold change) compared with WT adult RPW was significantly lower (*p*-value of 0.001) (Figure 4G). Similarly, the RferWhite homozygous mutant (−/−) carrying a 3-nt indel exhibited significantly lower relative expression levels (fold change) compared with WT adult (*p*-value of 0.001) (Figure 4F).

To comprehensively characterize the effect of the 13-nt CRISPR/Cas9-induced deletion on *cardinal* mRNA processing, the full-length *cardinal* ORF was amplified from cDNA obtained from the homozygous (−/−) G3 line (Figure S4). Sanger sequencing of the full-length ORF confirmed the presence of the 13-nt deletion (TGATGGTTGTAAC) in all homozygous mutant transcripts. The persistent translucent reddish-brown eye-color phenotype observed in all *cardinal* −/− adults is consistent with a complete loss of *cardinal* enzyme function and disruption of ommochrome pigment biosynthesis.

### Mating fecundity and X-linked inheritance of *white* and *cardinal* mutants

To evaluate mating performance and reproductive output under different mating ratios, a single RferWhite homozygous mutant female (w−/w−) was crossed with five WT males (W^+^/Y) in a mating chamber. These crosses produced an average of 54 eggs per female (n=3) over two weeks of collection. Similarly, crosses between a homozygous RferCar mutant female (c−/c−) and five WT males (C^+^/Y) laid an average of 47 eggs over a 2-week collection period. In contrast, control crosses of a WT female (W^+^/W^+^) with five WT males (W^+^/Y) produced 86 eggs. Reciprocal mating-ratio experiments were also performed. Five RferWhite homozygous mutant females (w−/w−) crossed with a single WT male (W^+^/Y) produced an average of 51 eggs/female over two weeks. Similarly, five RferCar homozygous mutant females (c−/c−) crossed with a single WT male (C^+^/Y) yielded an average of 54 eggs per female over the same period. Control single WT male (C^+^/Y) crossed with five WT females (C^+^/C^+^) produced an average of 58 eggs per female. All data were averages of three independent crosses. These results indicate no significant difference in fecundity between mutants and WT.

We determined the inheritance patterns of *white* and *cardinal* in *R. ferrugineus* by genotyping individual progeny *via* PCR amplification and Sanger sequencing of genomic regions flanking the sgRNA target sites. Both *white* and *cardinal* showed a classical X-linked inheritance pattern (Figure 5). A homozygous mutant RferWhite female (w−/w−) with a 9-nt deletion crossed with a WT male (W^+^/Y) produced progeny of heterozygous females (W^+^/w−) and hemizygous males (w−/Y) in an expected 1:1 ratio. Sequencing confirmed that all male offspring carried a 9-nt deletion (Figure 6A and B). Conversely, a hemizygous mutant RferWhite male (w−/Y) (9-nt deletion) crossed with a WT female (W^+^/W^+^) produced progeny of heterozygous females (W^+^/w−) and WT males (W^+^/Y) in the expected 1:1 ratio. All male offspring were WT, whereas all female offspring were heterozygous, as confirmed by Sanger sequencing (Figure 6C and D).

**Figure 5.**
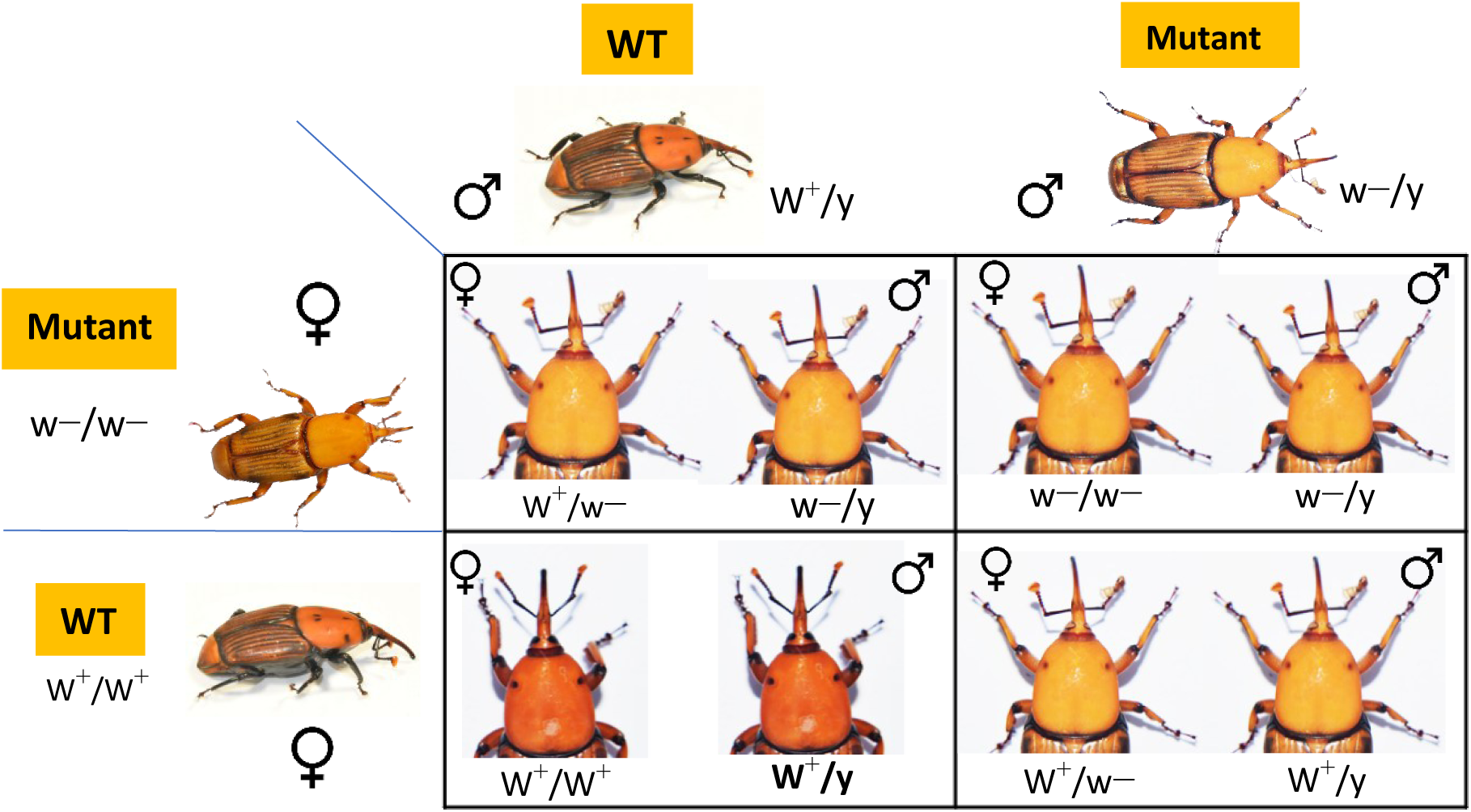
The X-linked inheritance patterns of RferWhite in *R. ferrugineus*. Germline-based homozygous (9-nt deletion) (w−/w−) were generated through the aforesaid crossing experiment depicted in Figure 4A and were verified through Sanger sequencing. The hemizygous line (9-nt deletion) male (w−/Y) or homozygous female (w−/w−) was crossed with wild-type female (W+/W+) or male (W+/Y), and the progeny genotype and phenotype are shown. The progeny’s genotype was verified by Sanger sequencing (see Figure 6). As the RfeCar crossing scheme (13-nt deletion) showed a similar X-linked inheritance pattern to RferWhite, the results are not shown.

**Figure 6.**
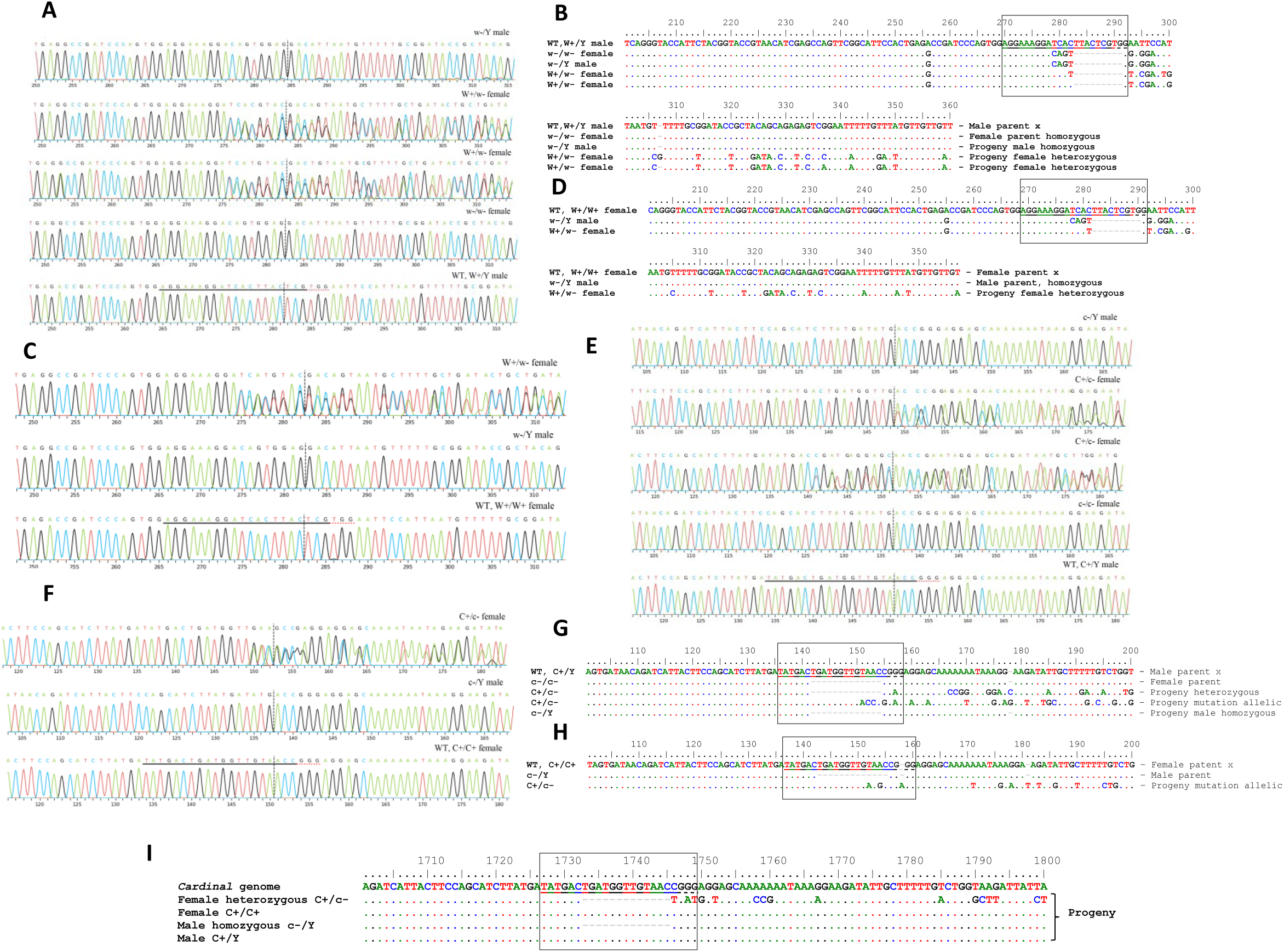
X-linked inheritance pattern of *white* and *cardinal* in *R. ferrugineus*. Two reciprocal crosses: homozygous mutant female (−/−) × wild-type (WT) male (+/Y), and hemizygous mutant male (−/Y) × WT female (+/+). For RferWhite and RferCar, homozygous mutants carrying 9-nt and 13-nt deletions, respectively, were selected. The Sanger sequence chromatogram alignment of parental and progeny of RferWhite (**A** and **C**) and RferCar (**E** and **F**) mutants compared with WT. This alignment of base calls from both the WT and mutant samples in the AB1 files was visualized using ICE *v*4 https://ice.editco.bio/. The horizontal, black-underlined region represents the guide sequence, and the red-dotted underline indicates the PAM site. The vertical black dotted line represents the actual cut site. **B.** Multiple sequence alignment of WT (+/Y) male x 9-nt RferWhite homozygous mutant female (−/−) and their progeny is shown. **D**. Multiple sequence alignment of WT female (+/+) x 9-nt RferWhite hemizygous mutant male (−/Y) and their progeny is shown. The gRNA target region is highlighted in a square box. **G.** Multiple sequence alignment of WT male (+/Y) x 13-nt indel RferCar homozygous (−/−) mutant female and their progeny is shown. **H**. Multiple sequence alignment of WT female x 13-nt indel RferCar hemizygous (−/Y) mutant male and their progeny is shown. The region corresponding to the guide RNA (underlined) and the PAM site (dotted line) is shown. **I**. Multiple sequence alignment of the progeny genotype from the WT male (+/Y) x 13-nt RferCar heterozygous (+/−) mutant female is shown. The number of deleted nucleotide sequences is shown as a grey dashed line.

The progeny of *RferCar* exhibited a similar X-linked inheritance pattern (Figure 5). A homozygous mutant RferCar female (c−/c−) crossed with a WT male (C^+^/Y) produced a progeny of heterozygous females (C^+^/c−) and hemizygous males (c−/Y) at the expected 1:1 Mendelian ratio. Sanger sequencing confirmed that all males carried the CRISPR-induced indel (Figure 6E and G). In contrast, a hemizygous RferCar mutant male (c−/Y) crossed with a WT female (C^+^/C^+^) produced heterozygous female (C^+^/c−) and WT male offspring (C^+^/Y) at the expected 1:1 ratio, which were later confirmed through Sanger sequencing (Figure 6F and H).

To further validate the inheritance pattern of *cardinal* in *R. ferrugineus*, the heterozygous females (C^+^/c−) carrying a 13-nt deletion were crossed with WT males (C^+^/Y). This cross produced progeny of heterozygous females (C^+^/c−) and WT females (C^+^/C^+^) at a 1:1 ratio, as well as WT males (C^+^/Y) and hemizygous males (c−/Y) at a 1:1 ratio (Figure 6I). All mutant males exhibited the characteristic translucent reddish-brown eye phenotype, and their genotype (c−/Y) was confirmed by Sanger sequencing (Figure 6I).

We performed an *in silico* MaxEntScan analysis for RferCar (Yeo and Burge, 2003) and predicted that the 13-nt deletion abolishes a strong canonical splice donor site (9-mer: ATG|GTTGTA; MaxEntScan score = 4.576) and six overlapping exonic splicing enhancer (ESE) hexamers (ACTGAT, CTGATG, TGATGG, GATGGT, ATGGTT, GGTTGT, reducing splice donor efficiency and potentially producing the minority intron-retained transcript population. In contrast, we screened the *cardinal* 13-nt mutant (−/−) ORF and found that the majority of transcripts carrying the 13-nt frameshift deletion were correctly spliced (Figure S4), which was predicted to introduce a premature termination codon (PTC) and generate a truncated, non-functional *cardinal* protein. Further, the persistent translucent reddish-brown eye phenotype in all *cardinal* (−/−) 13-nt deletion homozygous male adults is consistent with complete loss of *cardinal* function and disruption of ommochrome pigment biosynthesis.

## Discussion

Here, we establish the first CRISPR/Cas9-based genome editing platform for the globally invasive quarantine pest, the red palm weevil (*R. ferrugineus*). Specifically, we show that CRISPR/Cas9-mediated targeted disruption of two key factors in the ommochrome biosynthetic pathway — the ABC transporter *white*, which transports the intermediate precursor 3-hydroxykynurenine into pigment granules, and the heme peroxidase *cardinal*, which catalyzes the oxidative condensation of 3-hydroxykynurenine into ommochrome pigments — produced distinct, heritable eye-color phenotypes in *R. ferrugineus*. Disruption of *white* and *cardinal* produced a white-eyed phenotype with a brownish outer margin. In addition, the *cardinal* mutant produced translucent brownish-white eyes with white streaks, both arising from failures in ommochrome synthesis and deposition. We further demonstrate that *white* and *cardinal* deficiencies result in a persistent, heritable translucent reddish-brown eye phenotype in homozygous knockout adults. *R. ferrugineus* lines carrying a 9-nt deletion and a 13-nt frameshift-inducing deletion that generate premature termination codons and truncated, non-functional proteins were demonstrated, and both *white* and *cardinal* follow X-linked inheritance. CRISPR/Cas9-based genome editing has been applied to several insect species (Komal et al., 2023; Li et al., 2025; Singh et al., 2022; Sun et al., 2017); however, to our knowledge, CRISPR/Cas9-mediated genetic manipulation has not previously been achieved in the genus *Rhynchophorus* (Hoddle et al., 2026), nor in any true weevil (Curculionidae). Given the devastating impact of *R. ferrugineus* on global palm agriculture and cultural landscapes, targeting genes governing pivotal biological functions offers a promising foundation for developing genetics-based pest management strategies (Hoddle et al., 2024). The present study provides a detailed methodological framework for CRISPR/Cas9 genome editing in this weevil.

Coleoptera is the largest insect order (∼400,000 species), yet only eight species have confirmed CRISPR/Cas9 genome editing as of mid-2026 (Chu et al., 2018; Gui et al., 2020; Liu et al., 2025; Shirai and Daimon, 2020), reflecting the major challenges associated with embryo accessibility and limited genomic resources outside *Tribolium castaneum* (Arjunan and Thiruvengadam, 2026; Li et al., 2025; Saini et al., 2026). Palm weevils belong to Curculionidae, the largest beetle family with approximately 83,000 described species (Bandeira et al., 2021); however, prior to the present study, no genome-editing work on any Curculionid insect has been reported (Li et al., 2025). Genome editing in Coleoptera is complicated by the requirement for embryo microinjection during early embryogenesis, restricting CRISPR/Cas9-based targeted mutagenesis to a limited number of coleopteran species (Komal et al., 2023; Li et al., 2025; Singh et al., 2022). Furthermore, compared with other insect orders, the relative scarcity of genomic, transcriptomic, and functional resources for Coleoptera creates a significant knowledge gap that impedes the development of CRISPR-based gene editing. High-quality, well-annotated genome assemblies are an essential prerequisite for functional genomics, as minor variations in protein-coding sequences can compromise both protein function and the efficiency of RNA-guided editing. The availability of a chromosomal-level *R. ferrugineus* genome assembly (Sudalaimuthuasari et al., 2024) has enabled precise annotation of the *white* and *cardinal* genes. The quality of the Cas9 nuclease is equally critical for efficient genome editing; accordingly, we employed a HiFi *Streptococcus pyogenes* (S.p.) Cas9 nuclease, which minimizes off-target cleavage while maintaining robust on-target activity.

The ommochrome biosynthetic pathway is well characterized in insects, and eye pigmentation is primarily determined by ommochrome pigments (Ryall and Howells, 1974; Stavenga, 2002; Summers et al., 1982). Two terminal components, the ABC transporter *white* and the heme peroxidase *cardinal,* are essential for ommochrome synthesis. Functional disruption of either factor results in distinct eye-color phenotypes (Adrianos et al., 2018; Grubbs et al., 2015; Shirai et al., 2022). Phylogenetic analysis of *cardinal* and *white* within the broader context of the ommochrome biosynthesis pathway revealed evolutionary conservation, suggesting that the regulation of these eye-color genes has diversified across insect orders but is conserved within families and subfamilies. In the present study, both *white* and *cardinal* genes clustered within the same Curculionidae clade, establishing them as well-suited targets for initial CRISPR/Cas9 trials in this family and in Coleoptera more broadly, for demonstrating targeted gene disruption, germline inheritance, and progeny screening—particularly given their X-linked inheritance.

Differential expression analysis confirmed that both *white* and *cardinal* are highly expressed in the gut and head of *R. ferrugineus*, consistent with the established role of the ommochrome pathway in these tissues. However, qRT-PCR also revealed expression of both genes in wings and legs, consistent with the dual physiological roles of the ommochrome pathway in sensory and appendage tissues: detoxification of excess tryptophan and photoprotection of photoreceptors (Figon and Casas, 2019; How et al., 2023). Excess tryptophan is cytotoxic; the ommochrome pathway converts it into stable, insoluble pigments such as xanthommatin, which are sequestered in storage pigment granules, thereby protecting delicate neural and sensory tissues from oxidative damage. The resulting pigments also attenuate ultraviolet radiation and mitigate oxidative stress, supporting the high energetic demands of arthropod vision and mechanosensation (Figon and Casas, 2019). A change in whole-body coloration from reddish-brown to pale yellow was observed in *white* mutant *R. ferrugineus*, indicating that *white* disruption affects not only eye pigmentation but also overall body coloration. Insect body color is primarily governed by three core biochemical pathways — melanin, ommochrome, and pteridine (Bai et al., 2026) — and disruption of the ommochrome pathway likely accounts for the body color phenotype observed in *white* mutant *R. ferrugineus*. In appendage tissues, ommochrome precursors including 3-hydroxykynurenine and ommochrome pigments contribute to cuticular tanning and regional color patterning (How et al., 2023), and transcription factors have been shown to co-regulate both the melanin and ommochrome pathways during appendage development (Bayala et al., 2023). Notably, no apparent body color change was observed in *cardinal* mutant adults beyond the eye-color phenotype, suggesting that the role of *cardinal* in body coloration may be compensated by residual xanthommatin produced through spontaneous autoxidation of 3-hydroxykynurenine.

Transcript abundance of both *cardinal* and *white* was significantly reduced in homozygous mutant *R. ferrugineus* (Figure 4), and both phenotypic and genotypic analyses confirmed successful gene knockout. CRISPR/Cas9-induced double-strand breaks repaired by non-homologous end joining (NHEJ) generate insertions or deletions at the cut site, which frequently result in frameshift mutations and premature termination codons (Ran et al., 2013). RNA transcripts containing premature termination codons produce truncated coding sequences that interfere with protein synthesis and are subject to degradation by the nonsense-mediated mRNA decay (NMD) surveillance pathway (Karousis et al., 2016; Popp and Maquat, 2016), which may contribute to the observed reduction in transcript abundance. Full-length ORF sequencing and analysis of the gRNA-flanking region together provide a mechanistically complete account of the mRNA consequences of the CRISPR/Cas9-induced 13-nt deletion in *cardinal*. Full-length ORF amplification from cDNA confirmed that *cardinal* transcripts are correctly spliced in the homozygous (−/−) G3 line and carry the 13-nt frameshift deletion (TGATGGTTGTAAC) (Figure S4). *In silico* MaxEntScan analysis (Yeo and Burge, 2003) further predicted that the 13-nt deletion abolishes a strong canonical splice donor site (MaxEntScan score = 4.576; 9-mer: ATG|GTTGTA) and six overlapping exonic splicing enhancer (ESE) hexamers within the deleted sequence, thereby reducing spliceosome engagement efficiency at the adjacent intron. No new GT dinucleotide splice donor context was generated at the deletion junction (new junction: ATATG|ACCGG). However, sequencing of the full-length *cardinal* ORF from cDNA confirmed the presence of a correctly spliced transcript carrying the 13-nt deletion without intron retention (Figure S4), indicating that the MaxEntScan prediction of splice donor disruption did not alter the functional outcome: the correctly spliced transcript carries the 13-nt frameshift deletion, introducing a premature termination codon that abolishes *cardinal* enzyme activity (Popp and Maquat, 2016). The persistent and heritable translucent reddish-brown eye phenotype observed in all *cardinal* −/− G3 adults provides phenotypic evidence for complete functional loss of *cardinal* activity and disruption of ommochrome biosynthesis.

The *R. ferrugineus* karyotype comprises ten autosomes and one pair of sex chromosomes (X and Yp), with males being XY and females XX (Al-Qahtani et al., 2014), a system further supported by the near-chromosomal-level genome assembly generated using PacBio HiFi sequencing and Dovetail Omni-C reads (∼779 Mb; N50 ∼43 Mb; 99.5% BUSCO completeness) (Sudalaimuthuasari et al., 2024). No published, independently verified functional mapping of *white* or *cardinal* to the X chromosome in *R. ferrugineus* or any other Coleopteran species has been reported prior to this study. The present study provides the first functional evidence that both *white* and *cardinal* exhibit classical X-linked Mendelian inheritance in *R. ferrugineus*, based on systematic crossing experiments. Crosses between hemizygous mutant males and WT females produced exclusively heterozygous female progeny (+/−) displaying an intermediate eye-color phenotype, while crosses between homozygous mutant females and WT males produced both heterozygous female (+/−) and hemizygous mutant male (−/Y) progeny — a segregation pattern diagnostic of X-linked inheritance. The *white* gene is classically X-linked in *Drosophila,* but this pattern is not universal across insect orders (Jeffries et al., 2021). Comparative genomic analyses have established that the X chromosome in Coleoptera carries a distinct gene complement from that of Diptera, reflecting independent sex chromosome evolution across insect orders (Dutrillaux and Dutrillaux, 2017; Jeffries et al., 2021; Vicoso and Bachtrog, 2015), so X-linkage of *white* in *Drosophila* cannot be extrapolated to predict X-linkage in *R. ferrugineus*. Whether *white* or *cardinal* is X-linked in *T. castaneum* has not been independently confirmed (Grubbs et al., 2015). Genotypic screening of progeny from a *cardinal* 13-nt heterozygous female (+/−) crossed with a WT male (+/+) produced hemizygous mutant male progeny (−/Y) with a distinct eye-color phenotype, further corroborating the X-linked inheritance of *cardinal* in *R. ferrugineus*.

Sanger sequencing of heterozygous progeny frequently revealed overlapping chromatogram peaks at or immediately downstream of the cleavage site, or upstream of the PAM region—a well-recognized signature of heterozygosity following CRISPR/Cas9 editing, in which one chromosome carries the indel and the other the WT allele, producing superimposed signal traces. A subset of heterozygous (+/−) progeny exhibited mixed peaks across the entire amplified region (e.g., the 618 bp *cardinal* flanking amplicon). Deconvolution using the Synthego ICE online tool, followed by TA cloning and individual colony sequencing, confirmed that these samples carried one edited allele and one WT allele—with the chromatogram signal overlap arising from the heterozygous genotype rather than mosaicism. In contrast, animals displaying whole-region mixed peaks likely present mosaics or chimeras, comprising a mixture of WT, heterozygous, and biallelically edited cells. Additionally, preferential amplification of the shorter WT-like product over the mutant allele during RT-PCR may contribute to the apparent underrepresentation of the mutant allele, particularly when mutant mRNA is at low abundance or partially degraded (Connelly & Pruett-Miller, 2019; Tuladhar et al., 2019). Similar observations have been reported in CRISPR/Cas9 editing studies in *Spodoptera exigua* (Vatanparast et al., 2024). In contrast, crosses from two homozygous knockout adults (−/−) produced progeny with 100 % edited alleles — confirmed by clean, single-peak Sanger chromatograms spanning the CRISPR cleavage site — establishing a stable, genotypically uniform homozygous line (Figure 6).

In the red palm weevil, newly eclosed *cardinal* mutant adults initially exhibit brownish-white eyes with visible white streaks—affecting both eyes or either eye alone—which gradually transition to a persistent translucent reddish-brown eye color. This progressive color change is most parsimoniously explained by the spontaneous autoxidation of accumulated 3-hydroxykynurenine into xanthommatin over time (Osanai-Futahashi et al., 2016; Zhang et al., 2017), a phenomenon also documented in the *T. castaneum cardinal* mutant (Shirai and Daimon, 2020). *Cardinal* catalyzes the terminal enzymatic step of ommochrome biosynthesis to produce xanthommatin enzymatically (Figure 3); therefore, its disruption abolishes the canonical pathway. However, slow spontaneous autoxidation of the accumulated 3-hydroxykynurenine substrate can generate low levels of xanthommatin over time, accounting for the gradual color shift to a reddish-brown eye phenotype. *White* mutant *R. ferrugineus* adults exhibited a yellowish body color compared with the reddish-brown coloration of WT adults (Figure 3). In contrast, *cardinal* mutant adults showed no apparent body color changes beyond the eye phenotype. This difference likely reflects the distinct functional roles of the two components: *white* transports 3-hydroxykynurenine into pigment granules across multiple tissues, so loss of *white* disrupts ommochrome deposition throughout the body, whereas *cardinal* acts at the terminal enzymatic step, and spontaneous autoxidation may partially compensate for its loss in non-ocular tissues. Although both genes are expressed during *R. ferrugineus* pupal stages, no visible body color change was detected in *white* knockout pupae, suggesting that the ommochrome pathway’s contribution to body coloration becomes phenotypically apparent only in adults.

Given escalating concerns over palm tree protection and the urgent need for strategies that reduce reliance on chemical insecticides (Hoddle et al., 2024), genetics-based pest management approaches—including genome engineering—offer sustainable, species-specific alternatives, while also raising important biosecurity considerations. The cryptic larval stages of *Rhynchophorus* spp., which are concealed within palm trunks, render conventional insecticide-based control inherently ineffective. Over the past two decades, functional genomics studies in *R. ferrugineus* have characterised key biological processes including chemoreception, detoxification, digestion, reproduction, neurobiology, and immunity (Al-Ayedh, 2008; Antony et al., 2019; Antony et al., 2018; Antony et al., 2021; Antony et al., 2024; Rasool et al., 2021; Sattar et al., 2024; Soffan et al., 2016), establishing *R. ferrugineus* as an emerging model for developing genetic control technologies, including gene drive systems that may offer a sustainable alternative to conventional chemical management. The present study establishes a CRISPR/Cas9 genome-editing platform in *R. ferrugineus*, laying the groundwork for the accelerated exploration of genetic engineering approaches for sustainable palm weevil population management.

## Conclusion

This study demonstrates CRISPR/Cas9-mediated genome editing in the globally important invasive pest, the red palm weevil, *Rhynchophorus ferrugineus*, achieving functional disruption of two key components of the ommochrome biosynthetic pathway—the ABC transporter *white* and the heme peroxidase *cardinal*—thereby producing stable, heritable eye-color mutant lines. Critically, to our knowledge, this represents the first report of CRISPR/Cas9-mediated genome editing in any species of the family Curculionidae and in the quarantine pest genus *Rhynchophorus*. Our findings establish X-linked inheritance of both *white* and *cardinal* in *R. ferrugineus* and provide a systematic methodological framework for embryonic microinjection, mutant screening, and crossing strategies to generate homozygous knockout lines in Coleoptera. Beyond their immediate utility as visible genetic markers, *white* and *cardinal* represent valuable reporter genes for tracking genetic transformation and future gene drive experiments in *R. ferrugineus*, offering a tractable, visually screenable platform for future development of genetics-based biocontrol programmes for sustainable red palm weevil management.

## Acknowledgments

The authors extend their appreciation to the UAE Ministry of Climate Change and Environment for funding the red palm weevil research program. This work was supported by the Khalifa Center for Genetic Engineering and Biotechnology, UAE University, through grant no. KCGEB2. We thank Andrin Avery L. Tayde and Jeffry L. Pactao-in for their technical assistance with rearing red palm weevils, and Asma Al Bloushi and Shaika Al Amiri for their coordination and assistance with laboratory and administrative activities.

## Author contributions

K.A., B.A., and L.L planned and supervised the experiment. H.B.H., G.A.B., and B.A performed laboratory experiments. B.A. and J.J. contributed to the transcriptome analysis. B.A. wrote the paper. L.L and K.A reviewed and edited the paper, and all authors approved the final version of the manuscript.

## Availability of data and materials

The mutant red palm weevil *white* and *cardinal* homozygous lines are maintained at the insect rearing facility at the Khalifa Center for Genetic Engineering and Biotechnology (KCGEB).

## Supplementary data

*Supplementary data to this article can be found online at:*

**Table S1.** The semi-synthetic diet was used to rear the red palm weevil (*R. ferrugineus*).

**Table S2.** List of the oligonucleotide primers used in the current study.

**Table S3** NCBI accession numbers for different body parts of the Red Palm Weevil (*Rhynchophorus ferrugineus*). Data are collected from male and female specimens under both laboratory and field conditions, as reported in the SRA.

**Table S3a:** Red palm weevil antennal transcriptome assembly report.

**Table S3b:** Red palm weevil snout transcriptome assembly report.

**Table S3c:** Red palm weevil head transcriptome assembly report.

**Table S3d:** Red palm weevil leg transcriptome assembly report.

**Table S3e:** Red palm weevil wing transcriptome assembly report.

**Table S3f:** Red palm weevil thorax transcriptome assembly report.

**Table S3g**. Red palm weevil gut transcriptome assembly report.

**Table S3h:** Red palm weevil abdomen transcriptome assembly report.

**Figure S1** RferWhite Open Reading Frame (ORF). The region corresponding to the guide RNA (underlined) and the PAM site (dotted line) are shown.

**Figure S1a:** *R. ferrugineus white* (RferWhite) protein motif and domain profile.

**Figure S2** RferCar Open Reading Frame (ORF). The region corresponding to the guide RNA (underlined) and the PAM site (dotted line) are shown.

**Figure S2a:** *R. ferrugineus cardinal* (RferCar) protein motif and domain profile.

**Figure S3**. RferWhite and RferCar tissue-specific expression gel images.

**Figure S4.** Primer walking (using RfCar-GT-R1) (Table S2) Sanger sequence chromatogram (anti-sense strand) of RferCar ORF region corresponding to the guide RNA of *cardinal* mutant 13-nt homozygous RPWs. The arrow indicates the deleted ‘GTTACAACCATCA’ sequence from the gRNA region and the PAM shown in the box.

## References

1. Adrianos, S., Lorenzen, M., Oppert, B., 2018. Metabolic pathway interruption: CRISPR/Cas9-mediated knockout of tryptophan 2, 3-dioxygenase in Tribolium castaneum. Journal of insect physiology 107, 104–109.

2. Al-Ayedh, H., 2008. Evaluation of date palm cultivars for rearing the red date palm weevil, Rhynchophorus ferrugineus (Coleoptera: Curculionidae). Florida Entomologist 91, 353–358.

3. Al-Qahtani, A.H., Al-Khalifa, M.S., Al-Saleh, A.A., 2014. Karyotype, meiosis and sperm formation in the red palm weevil Rhynchophorus ferrugineus. Cytologia 79, 235–242.

4. Antony, B., Johny, J., Abdelazim, M.M., Jakše, J., Al-Saleh, M.A., Pain, A., 2019. Global transcriptome profiling and functional analysis reveal that tissue-specific constitutive overexpression of cytochrome P450s confers tolerance to imidacloprid in palm weevils in date palm fields. BMC genomics 20, 440.

5. Antony, B., Johny, J., Aldosari, S.A., 2018. Silencing the odorant binding protein RferOBP1768 reduces the strong preference of palm weevil for the major aggregation pheromone compound ferrugineol. Frontiers in physiology 9, 252.

6. Antony, B., Johny, J., Montagné, N., Jacquin-Joly, E., Capoduro, R., Cali, K., Persaud, K., Al-Saleh, M.A., Pain, A., 2021. Pheromone receptor of the globally invasive quarantine pest of the palm tree, the red palm weevil (Rhynchophorus ferrugineus). Molecular ecology 30, 2025–2039.

7. Antony, B., Montagné, N., Comte, A., Mfarrej, S., Jakše, J., Capoduro, R., Shelke, R., Cali, K., AlSaleh, M.A., Persaud, K., Pain, A., Jacquin-Joly, E., 2024. Deorphanizing an odorant receptor tuned to palm tree volatile esters in the Asian palm weevil sheds light on the mechanisms of palm tree selection. Insect Biochemistry and Molecular Biology, 104129.

8. Antony, B., Soffan, A., Jakše, J., Abdelazim, M.M., Aldosari, S.A., Aldawood, A.S., Pain, A., 2016. Identification of the genes involved in odorant reception and detection in the palm weevil Rhynchophorus ferrugineus, an important quarantine pest, by antennal transcriptome analysis. BMC genomics 17, 1–22.

9. Arjunan, N.K., Thiruvengadam, V., 2026. Harnessing CRISPR-Cas technology for insect pest control: current advances and future perspectives. Molecular Biology Reports 53, 856.

10. Bandeira, P.T., Fávaro, C.F., Francke, W., Bergmann, J., Zarbin, P.H.G., 2021. Aggregation pheromones of weevils (Coleoptera: Curculionidae): advances in the identification and potential uses in semiochemical-based pest management strategies. Journal of Chemical Ecology 47, 968–986.

11. Bayala, E.X., VanKuren, N., Massardo, D., Kronforst, M., 2023. aristaless1 has a dual role in appendage formation and wing color specification during butterfly development. BMC biology 21, 100.

12. Chu, F.-C., Wu, P.-S., Pinzi, S., Grubbs, N., Lorenzen, M.D.J.J.o.V.E.J., 2018. Microinjection of western corn rootworm, Diabrotica virgifera virgifera, embryos for germline transformation, or CRISPR/Cas9 genome editing. 57497.

13. Dress, A.W., Flamm, C., Fritzsch, G., Grünewald, S., Kruspe, M., Prohaska, S.J., Stadler, P.F., 2008. Noisy: identification of problematic columns in multiple sequence alignments. Algorithms for Molecular Biology 3, 7.

14. Dutrillaux, A.-M., Dutrillaux, B., 2017. Evolution of the sex chromosomes in beetles. I. The loss of the Y chromosome. Cytogenetic Genome Research 152, 97–104.

15. Figon, F., Casas, J., 2019. Ommochromes in invertebrates: biochemistry and cell biology. Biological Reviews 94, 156–183.

16. Fu, L., Li, P., Rui, Z., Sun, J., Yang, J., Wang, Y., Jia, D., Hu, J., Li, X., Ma, R., 2025. CRISPR/Cas9-Mediated Knockout of the White Gene in Agasicles hygrophila. International Journal of Molecular Sciences 26, 4586.

17. Gonzalez, F., Johny, J., Walker III, W.B., Guan, Q., Mfarrej, S., Jakše, J., Montagne, N., Jacquin-Joly, E., Alqarni, A.S., Al-Saleh, M.A., Pain, A., Antony, B., 2021. Antennal transcriptome sequencing and identification of candidate chemoreceptor proteins from an invasive pest, the American palm weevil, Rhynchophorus palmarum. Scientific reports 11, 8334.

18. Grubbs, N., Haas, S., Beeman, R.W., Lorenzen, M.D., 2015. The ABCs of eye color in Tribolium castaneum: orthologs of the Drosophila white, scarlet, and brown genes. Genetics 199, 749–759.

19. Gui, S., Taning, C.N.T., Wei, D., Smagghe, G.J.J.o.i.p., 2020. First report on CRISPR/Cas9-targeted mutagenesis in the Colorado potato beetle, Leptinotarsa decemlineata. 121, 104013.

20. Hoddle, M., Antony, B., El-Shafie, H., Chamorro, L., Milosavljević, I., B, L., Faleiro, R., 2024. Taxonomy, Biology, Symbionts, Omics, and Management of Rhynchophorus Palm Weevils (Coleoptera: Curculionidae: Dryophthorinae). Annual Review of Entomology 69, 449–479.

21. Hoddle, M.S., Antony, B., Torres, J., San Jose, G., Milosavljević, I., Hoddle, C.D., 2026. Taxonomy, biology, invasion history, and management of Rhynchophorus palmarum (Coleoptera: Curculionidae: Dryophthorinae). Journal of Integrated Pest Management 17, pmag016.

22. How, S.H.C., Banerjee, T.D., Monteiro, A., 2023. Vermilion and cinnabar are involved in ommochrome pigment biosynthesis in eyes but not wings of Bicyclus anynana butterflies. Scientific Reports 13, 9368.

23. Jeffries, D.L., Gerchen, J.F., Scharmann, M., Pannell, J.R., 2021. A neutral model for the loss of recombination on sex chromosomes. Philosophical Transactions of the Royal Society B: Biological Sciences 376, 20200096.

24. Karousis, E.D., Nasif, S., Mühlemann, O., 2016. Nonsense-mediated mRNA decay: novel mechanistic insights and biological impact. Wiley Interdisciplinary Reviews: RNA 7, 661–682.

25. Komal, J., Desai, H., Samal, I., Mastinu, A., Patel, R., Kumar, P.D., Majhi, P.K., Mahanta, D.K., Bhoi, T.K., 2023. Unveiling the genetic symphony: Harnessing CRISPR-Cas genome editing for effective insect pest management. Plants 12, 3961.

26. Lemoine, F., Correia, D., Lefort, V., Doppelt-Azeroual, O., Mareuil, F., Cohen-Boulakia, S., Gascuel, O., 2019. NGPhylogeny.fr: New generation phylogenetic services for non-specialists. Nucleic acids research 47, W260–265.

27. Li, J., Wang, J., Taning, C.N., De Schutter, K., 2025. Genetic engineering and genetic control in Coleoptera: progress, challenges and perspectives. Entomologia Generalis 45, 1591–1604.

28. Liu, Z.L., Zhou, Y.Y., Xu, Q.X., Wang, X.C., Liu, T.X., Tian, H.G.J.I.S., 2025. Efficient CRISPR-mediated genome editing can be initiated by embryonic injection but not by ovarian delivery in the beetle Tribolium castaneum. 32, 1185–1200.

29. Livak, K.J., Schmittgen, T.D., 2001. Analysis of relative gene expression data using real-time quantitative PCR and the 2− ΔΔCT method. Methods 25, 402–408.

30. Nguyen, L.T., Schmidt, H.A., Von Haeseler, A., Minh, B.Q., 2015. IQ-TREE: A fast and effective stochastic algorithm for estimating maximum-likelihood phylogenies. Molecular Biology and Evolution 32, 268–274.

31. Osanai-Futahashi, M., Tatematsu, K., Futahashi, R., Narukawa, J., Takasu, Y., Kayukawa, T., Shinoda, T., Ishige, T., Yajima, S., Tamura, T., 2016. Positional cloning of a Bombyx pink-eyed white egg locus reveals the major role of cardinal in ommochrome synthesis. Heredity 116, 135–145.

32. Popp, M.W., Maquat, L.E., 2016. Leveraging rules of nonsense-mediated mRNA decay for genome engineering and personalized medicine. Cell 165, 1319–1322.

33. Ran, F.A., Hsu, P.D., Wright, J., Agarwala, V., Scott, D.A., Zhang, F., 2013. Genome engineering using the CRISPR-Cas9 system. Nature protocols 8, 2281–2308.

34. Rasool, K.G., Mehmood, K., Tufail, M., Husain, M., Alwaneen, W.S., Aldawood, A.S., 2021. Silencing of vitellogenin gene contributes to the promise of controlling red palm weevil, Rhynchophorus ferrugineus (Olivier). Scientific Reports 11, 21695.

35. Ryall, R.L., Howells, A., 1974. Ommochrome biosynthetic pathway of Drosophila melanogaster: Variations in levels of enzyme activities and intermediates during adult development. Insect Biochemistry 4, 47–61.

36. Saini, A., Sharma, N., Sharma, N., Kumari, N., Sharma, M., Singh, B., Thakur, A.K., 2026. Precision pest management: Genome editing tools, specifically CRISPR/Cas9 and future prospects. Pesticide biochemistry and physiology, 106941.

37. Sattar, M.N., Naqqash, M.N., Rezk, A.A., Mehmood, K., Bakhsh, A., Elshafie, H., Al-Khayri, J.M., 2024. Sprayable RNAi for silencing of important genes to manage red palm weevil, Rhynchophorus ferrugineus (Coleoptera: Curculionidae). PloS one 19, e0308613.

38. Scieuzo, C., Rinaldi, R., Giglio, F., Salvia, R., Ali AlSaleh, M., Jakše, J., Pain, A., Antony, B., Falabella, P., 2024. Identification of multifunctional putative bioactive peptides in the insect model red palm weevil (Rhynchophorus ferrugineus). Biomolecules 14, 1332.

39. Shamim, G., Ranjan, S.K., Pandey, D.M., RAmANI, R., 2014. Biochemistry and biosynthesis of insect pigments. European Journal of Entomology 111, 149–164.

40. Shirai, Y., Daimon, T., 2020. Mutations in cardinal are responsible for the red-1 and peach eye color mutants of the red flour beetle Tribolium castaneum. Biochemical Biophysical Research Communications 529, 372–378.

41. Shirai, Y., Piulachs, M.-D., Belles, X., Daimon, T., 2022. DIPA-CRISPR is a simple and accessible method for insect gene editing. Cell Reports Methods 2.

42. Singh, S., Rahangdale, S., Pandita, S., Saxena, G., Upadhyay, S.K., Mishra, G., Verma, P.C., 2022. CRISPR/Cas9 for insect pests management: a comprehensive review of advances and applications. Agriculture 12, 1896.

43. Soffan, A., Antony, B., Abdelazim, M., Shukla, P., Witjaksono, W., Aldosari, S.A., Aldawood, A.S., 2016. Silencing the olfactory co-receptor RferOrco reduces the response to pheromones in the red palm weevil, Rhynchophorus ferrugineus. PloS one 11, e0162203.

44. Stavenga, D., 2002. Colour in the eyes of insects. Journal of Comparative Physiology A 188, 337–348.

45. Sudalaimuthuasari, N., Kundu, B., Hazzouri, K.M., Amiri, K.M., 2024. Near-chromosomal-level genome of the red palm weevil (Rhynchophorus ferrugineus), a potential resource for genome-based pest control. Scientific data 11, 45.

46. Summers, K., Howells, A., Pyliotis, N., 1982. Biology of eye pigmentation in insects, Advances in insect physiology. Elsevier, pp. 119–166.

47. Sun, D., Guo, Z., Liu, Y., Zhang, Y., 2017. Progress and prospects of CRISPR/Cas systems in insects and other arthropods. Frontiers in physiology 8, 608.

48. Vatanparast, M., Esmaeily, M., Stanley, D., Kim, Y., 2024. A PLA2 deletion mutant using CRISPR/Cas9 coupled to RNASeq reveals insect immune genes associated with eicosanoid signaling. PloS one 19, e0304958.

49. Vicoso, B., Bachtrog, D., 2015. Numerous transitions of sex chromosomes in Diptera. PLoS biology 13, e1002078.

50. Yeo, G., Burge, C.B., 2003. Maximum entropy modeling of short sequence motifs with applications to RNA splicing signals, Proceedings of the seventh annual international conference on Research in computational molecular biology, pp. 322–331.

51. Zhang, H., Lin, Y., Shen, G., Tan, X., Lei, C., Long, W., Liu, H., Zhang, Y., Xu, Y., Wu, J., 2017. Pigmentary analysis of eggs of the silkworm Bombyx mori. Journal of insect physiology 101, 142–150.

