## Supplementary material for "Ommochrome pathway knockout via CRISPR/Cas9 reveals sex-linked eye pigmentation and establishes a heritable genome-editing platform in Rhynchophorus ferrugineus": Table S1, Table S2, Table S3, Figure S1, Figure S1a, Figure S2, Figure S2a, Figure S3 and Figure S4

**Table S1** The semi-synthetic diet was used to rear the red palm weevil (*R. ferrugineus*).

| Ingredients | Amount |
|---|---|
| Cornmeal | 312g |
| Yeast | 312g |
| Agar | 140g |
| Wheat germ | 312g |
| Boiled coconut fibre | 744g |
| Ascorbic acid | 50mL (15g dissolved in 50mL distilled water) |
| Methyl paraben | 60mL (7.5g dissolve in 50mL 100% ethanol) |
| Propionic acid | 24mL |
| Multivitamins | 12pcs |
| Water | 5L |

**Table S2** List of the oligonucleotide primers used in the current study

| | |
|---|---|
| RfCar-GT-F1 | GGACCAAGAGAACAGATGAATCA |
| RfCar-GT-R1 | CATTTGCAACGCTAGGGTCA |
| RfCar-GT-F2 | TGGCTCCGAAAGATTCAAACA |
| RfCar-GT-R2 | TGGCCTATAAGCAACTCCGT |
| RferCar-FL-F | ATGTCTCAAAGTCAATTTAATGAAA |
| RferCar-qPCR-F | AACAGCTACGGAGTTGCTTATAG |
| RferCar-qPCR-R | TACGGGCAGACGGTAGATTA |
| RferWhite-qPCR-F | ACATGTTCATCGGTACCCTTAC |
| RferWhite-qPCR-R | CACTTCTTCGACTCTGGCTATT |
| RferCar-FL-R1 | TTAATTAGCAGGTTTTTCATAA |
| RferWhite-FL-F | ATGAATAACGAGGAGAAAATACC |
| RferWhite-FL-R | TCATTGGTCCTTATAAGTTCTCCT |
| RferWhite-PW-F1 | GTACCATTCTACGGTACCG |
| RferWhite-PW-F2 | GAACACCTGACCTTTCAAGCC |
| RferWhite-PW-F3 | GGGAACCCCAGAAGAAGCGGA |
| RferWhite-PW-F4 | GCGTGAGCACAAAAATGGCATG |
| RferWhite-PW-F5 | CTATACCGATCTATTTCGA |
| RferTubulin_F | GCTACCTTCATCGGCAACTC |
| RferTubulin_R | CCTTCGCCAAGTGATATAG |
| Rferβ-actin_F | AAAGGTTCCGTTGCCCTGAA |
| Rferβ-actin_R | TGGCGTACAAGTCCTTCCTG |

**Table S3** NCBI Accession numbers for different body parts of the Red Palm Weevil (*Rhynchophorus ferrugineus*). Data are collected from male and female specimens under both laboratory and field conditions, as reported in the SRA**.**

| | SRA | | | |
|---|---|---|---|---|
| **Part of the body** | **Male Laboratory** | **Male Field** | **Female Laboratory** | **Female Field** |
| Abdomen | - | - | SRR27695093 | SRR27695094 |
| Antennae | SRR22098126 | SRR22098127 | SRR22098128 | SRR22098129 |
| Fat body | SRR29346546 | SRR29346547 | - | |
| Head | SRR29346787 | SRR29346788 | SRR29346789 | SRR29346790 |
| Gut | - | SRR27695095 | - | SRR27695096 |
| Legs | SRR29221206 | SRR29221207 | SRR29221208 | SRR29221209 |
| Snout | SRR17732026 | SRR17732027 | SRR17732028 | SRR17732029 |
| Thorax | SRR29346791 | SRR29346792 | SRR29346793 | SRR29346794 |
| Wings | SRR29346548 | SRR29346549 | SRR29346550 | SRR29346551 |

**Table S3a:** Red palm weevil antennal transcriptome assembly report.

| | Male field | Female field | Male lab | Female lab |
|---|---|---|---|---|
| Total number of raw reads | 99,146,450 | 134,645,400 | 90,215,866 | 198.956.016 |
| Total length of reads (bp) | 14,971,113,950 | 20,331,455,400 | 13,622,595,766 | 30.042.358.416 |
| Total number of cleaned reads | 98,868,937 | 134,414,781 | 90,122,367 | 198,763,141 |
| Total length of cleaned reads (bp) | 12,753,517,786 | 17,740,681,985 | 11,867,016,957 | 26,492,600,964 |
| Number of contigs | 53,645 | 59,627 | 50,519 | 81,862 |
| Total length | 31,354,749 | 35,944,586 | 30,782,692 | 45,247,906 |
| -N50 | 772 | 860 | 822 | 656 |
| Average | 584 | 603 | 609 | 553 |
| -Min | 74 | 104 | 85 | 101 |
| -Max | 21,051 | 25,370 | 14,929 | 21,384 |

**Table S3b:** Red palm weevil snout transcriptome assembly report.

| | Male field | Female field | Male lab | Female lab |
|---|---|---|---|---|
| Total number of raw reads | 133,189,166 | 254,584,320 | 102,484,022 | 200,404,720 |
| Total length of reads (bp) | 20,111,564,066 | 38,442,232,320 | 15,475,087,322 | 30,261,112,720 |
| Total number of reads cleaned | 133,129,330 | 254,460,904 | 102,403,197 | 200,277,342 |
| Total length of reads cleaned (bp) | 18,172,140,980 | 34,453,144,617 | 13,685,276,150 | 26,990,967,531 |
| Number of contigs | 126,191 | 63,128 | 62,641 | 120,850 |
| Total length | 64,605,550 | 37,133,236 | 37,424,798 | 69,131,965 |
| -N50 | 588 | 792 | 777 | 702 |
| Average | 512 | 588 | 597 | 572 |
| -Min | 104 | 87 | 110 | 72 |
| -Max | 34,191 | 30,694 | 13,256 | 21,750 |

**Table S3c:** Red palm weevil head transcriptome assembly report.

| | Male field | Female field | Male lab | Female lab |
|---|---|---|---|---|
| Total number of raw reads | 54,557,564 | 62,318,808 | 83,419,136 | 93,499,916 |
| Total length of reads (bp) | 8,238,192,164 | 9,410,140,008 | 12,596,289,536 | 14,118,487,316 |

| | | | | |
|---|---|---|---|---|
| Total number of reads cleaned | 54,536,099 | 62,306,572 | 83,397,136 | 93,470,704 |
| Total length of reads cleaned (bp) | 7,097,516,043 | 8,256,465,906 | 11,307,790,700 | 12,606,154,439 |
| Number of contigs | 30,748 | 28,195 | 35,527 | 35,232 |
| Total length | 18,758,492 | 21,255,540 | 27,520,948 | 26,144,688 |
| -N50 | | | | |
| Average | 848 | 710 | 699 | 682 |
| -Min | 53 | 3 | 48 | 107 |
| -Max | 13,320 | 22,558 | 12,533 | 16,119 |

**Table S3d:** Red palm weevil leg transcriptome assembly report.

| | **Male field** | **Female field** | **Male lab** | **Female lab** |
|---|---|---|---|---|
| Total number of raw reads | 129,529,916 | 166,095,222 | 213,480,874 | 87,163,312 |
| Total length of reads (bp) | 19,559,017,316 | 25,080,378,522 | 32,235,611,974 | 13,161,660,112 |
| Total number of reads cleaned | 129,483,077 | 166,025,634 | 213,341,091 | 86,953,254 |
| Total length of reads cleaned (bp) | 17,609,846,633 | 22,645,601,080 | 28,909,590,744 | 10,771,202,802 |
| Number of contigs | 75,700 | 68,240 | 105,309 | 70,207 |
| Total length | 40,416,179 | 37,689,152 | 61,652,451 | 37,541,225 |
| -N50 | 602 | 647 | 734 | 643 |
| Average | 515 | 528 | 585 | 524 |
| -Min | 29 | 61 | 102 | 29 |
| -Max | 23,454 | 25,849 | 23,303 | 12,319 |

**Table S3e:** Red palm weevil wing transcriptome assembly report.

| | **Male field** | **Female field** | **Male lab** | **Female lab** |
|---|---|---|---|---|
| Total number of raw reads | 57,223,934 | 75,845,290 | 97,665,294 | 116,092,070 |
| Total length of reads (bp) | 8,640,814,034 | 11,452,638,790 | 14,747,459,394 | 17,529,902,570 |
| Total number of reads cleaned | 57,199,080 | 75,824,164 | 97,627,819 | 116,038,284 |
| Total length of reads cleaned (bp) | 7,564,707,820 | 10,224,781,677 | 13,183,228,532 | 15,631,434,509 |

| | | | | |
|---|---|---|---|---|
| Number of contigs | 73,685 | 72,889 | 38,804 | 46,395 |
| Total length | 37,562,329 | 34,046,619 | 26,501,763 | 29,720,806 |
| -N50 | 586 | 516 | 990 | 881 |
| Average | 496 | 454 | 631 | 604 |
| -Min | 23 | 52 | 17 | 5 |
| -Max | 19,418 | 18,328 | 21,964 | 17,855 |

**Table S3f:** Red palm weevil thorax transcriptome assembly report.

| | **Male field** | **Female field** | **Male lab** | **Female lab** |
|---|---|---|---|---|
| Total number of raw reads | 127,276,228 | 38,248,088 | 45,162,240 | 66,814,704 |
| Total length of reads (bp) | 19,218,710,428 | 5,775,461,288 | 6,819,498,240 | 10,089,020,304 |
| Total number of reads cleaned | 127,178,496 | 38,215,277 | | 66,773,773 |
| Total length of reads cleaned (bp) | 16,835,875,428 | 5,102,840,160 | | 8,933,323,672 |
| Number of contigs | 30,708 | 24,289 | | 28,064 |
| Total length | 22,075,097 | 17,622,367 | | 20,591,404 |
| -N50 | 1,125 | 1,058 | | 1,058 |
| Average | 680 | 696 | | 688 |
| -Min | 16 | 52 | | 9 |
| -Max | 39,370 | 14,640 | | 17,209 |

**Table S3g**. Red palm weevil gut transcriptome assembly report.

| | Male gut SRR27695095 | Female gut SRR27695096 |
|---|---|---|
| Total number of raw reads | 26,779,316 | 215,865,354 |
| Total length of reads (bp) | 4,043,676,716 | 32,595,668,454 |
| Total number of reads cleaned | 26,765,101 (10,185,172 pairs, 3,218,761 singles) | 215,734,706 (76,235,113 pairs, 47,326,535 singles) |
| Total length of reads cleaned (bp) | 3,564,415,653 | 29,435,142,751 |
| Number of contigs | 31,312 | 67,747 |
| Total length | 24,182,277 | 44,910,876 |
| N50 | 1,305 | 1,018 |
| Average | 772 | 663 |
| -Min | 80 | 87 |
| -Max | 14,443 | 25,094 |

**Table S3h:** Red palm weevil abdomen transcriptome assembly report.

| | Female field | Female lab |
|---|---|---|
| Total number of raw reads | 67,353,892 | 44,502,316 |
| Total length of reads (bp) | 10,170,437,692 | 6,719,849,716 |
| Total number of reads cleaned | 44,469,373 | 67,264,457 |
| Total length of reads cleaned (bp) | 5,544,381,577 | 8,117,880,819 |
| Number of contigs | 27,961 | 24,564 |
| Total length | 21,005,763 | 13,470,612 |
| -N50 | 1,194 | 722 |
| Average | 713 | 532 |
| -Min | 93 | 47 |
| -Max | 8,383 | 8,417 |

**Figure S1** RferWhite Open Reading Frame (ORF). The region corresponding to the guide RNA (underlined) and the PAM site (dotted line) are shown.

```
                       10        20        30        40        50        60        70        80        90       100
              ....|....|....|....|....|....|....|....|....|....|....|....|....|....|....|....|....|....|....|....|
RferWhite_ORF ATGAATAACGAGGAGAAAATACCCTTGTTTTCAAGTAATGGTTACGGTGACGTGGACCAGGGTAATGATAATTCTTCATCGGAGAGTTCCGGTTCCTTCA
               M  N  N  E  E  K  I  P  L  F  S  S  N  G  Y  G  D  V  D  Q  G  N  D  N  S  S  S  E  S  S  G  S  F

                      110       120       130       140       150       160       170 gRNA1 180       190       200
              ....|....|....|....|....|....|....|....|....|....|....|....|....|....|....|....|....|....|....|....|
RferWhite_ORF GGGTACCATTCTACGGTACCGTAACATCGAGCCAGTTCGGCATTCCACTGAGACCGATCCCAGTGGAGGAAAGGATCACTTACTCGTGGAATTCCATTAA
              R  V  P  F  Y  G  T  V  T  S  S  Q  F  G  I  P  L  R  P  I  P  V  E  E  R  I  T  Y  S  W  N  S  I  N

                      210       220       230       240       250       260       270       280       290       300
              ....|....|....|....|....|....|....|....|....|....|....|....|....|....|....|....|....|....|....|....|
RferWhite_ORF TGTTTTTGCGGATACCGCTTCAGCAGAGAGTCGGAATTTTTGTTTATGTTGTTGTTCAAATGGCGAGTACGAAAATTTACAGAAAAAACATCTAGTGAAA
                V  F  A  D  T  A  S  A  E  S  R  N  F  C  L  C  C  C  S  N  G  E  Y  E  N  L  Q  K  K  H  L  V  K

                      310       320       330       340       350       360       370       380       390       400
              ....|....|....|....|....|....|....|....|....|....|....|....|....|....|....|....|....|....|....|....|
RferWhite_ORF AACGTGAACGGCGTGGCTCGTCCAGGCGAACTGCTGGCAATTTTAGGCTCAAGCGGAGCGGGCAAAACCACCCTCCTCAACGCGTTGACCTTTCGCTCGT
                N  V  N  G  V  A  R  P  G  E  L  L  A  I  L  G  S  S  G  A  G  K  T  T  L  L  N  A  L  T  F  R  S

                      410       420       430       440       450       460       470       480       490       500
              ....|....|....|....|....|....|....|....|....|....|....|....|....|....|....|....|....|....|....|....|
RferWhite_ORF CGAAGAACATCAGCGTGACGGGCACACGCTGCGTCAACGGCAACCCGGTACACTCGACGGCGCTCACCAGCATCTCGGCCTACGTCCAGCAAGCCGACAT
              S  K  N  I  S  V  T  G  T  R  C  V  N  G  N  P  V  H  S  T  A  L  T  S  I  S  A  Y  V  Q  Q  A  D  M

                      510       520       530       540       550       560       570       580       590       600
              ....|....|....|....|....|....|....|....|....|....|....|....|....|....|....|....|....|....|....|....|
RferWhite_ORF GTTCATCGGTACCCTTACCGTCCGGGAACACCTGACCTTTCAAGCCCTTGTTAGAATGGACAGACATCTCAAGTACGACCAACGAATAGCCAGAGTCGAA
               F  I  G  T  L  T  V  R  E  H  L  T  F  Q  A  L  V  R  M  D  R  H  L  K  Y  D  Q  R  I  A  R  V  E

                      610       620       630       640       650       660       670       680       690       700
              ....|....|....|....|....|....|....|....|....|....|....|....|....|....|....|....|....|....|....|....|
RferWhite_ORF GAAGTGATATCAGAGATGTCACTGAATAAATGCGAGAATACCTGTATAGGAGTGCCAGGAAGACTGAAAGGCCTGTCAGGTGGCGAGCAGAAGAGACTAT
               E  V  I  S  E  M  S  L  N  K  C  E  N  T  C  I  G  V  P  G  R  L  K  G  L  S  G  G  E  Q  K  R  L

                      710       720       730       740       750       760       770       780       790       800
              ....|....|....|....|....|....|....|....|....|....|....|....|....|....|....|....|....|....|....|....|
RferWhite_ORF CGTTCGCTTCAGAGATGCTTACCAACCCATCGCTCATGTTCTGCGACGAACCAACATCAGGTTTAGATTCCTTTATGGCACTCAACGTCGTACAAATCTT
              S  F  A  S  E  M  L  T  N  P  S  L  M  F  C  D  E  P  T  S  G  L  D  S  F  M  A  L  N  V  V  Q  I  L

                      810       820 gRNA2 830       840       850       860       870       880       890       900
              ....|....|....|....|....|....|....|....|....|....|....|....|....|....|....|....|....|....|....|....|
RferWhite_ORF GAAGCAAATGGCCCACACTGGTAAGACCGTAATCTGTACGATCCATCAGCCATCGTCCGAACTATACGCCATGTTCGATAAGCTGCTCCTGATGGCAGAA
                K  Q  M  A  H  T  G  K  T  V  I  C  T  I  H  Q  P  S  S  E  L  Y  A  M  F  D  K  L  L  L  M  A  E

                      910       920       930       940       950       960       970       980       990      1000
              ....|....|....|....|....|....|....|....|....|....|....|....|....|....|....|....|....|....|....|....|
RferWhite_ORF GGACGGATCGCCTTTCTGGGAACCCCAGAAGAAGCGGACATATTTTTCAGGGAATTAGATGCCCCATGTCCCAAGAATTATAATCCTGCCGATTATTTTA
```

```
                G  R  I  A  F  L  G  T  P  E  E  A  D  I  F  F  R  E  L  D  A  P  C  P  K  N  Y  N  P  A  D  Y  F

                       1010      1020      1030      1040      1050      1060      1070      1080      1090      1100
               ....|....|....|....|....|....|....|....|....|....|....|....|....|....|....|....|....|....|....|....|
RferWhite_ORF  TTCAGCTATTGGCAATAGTTCCGGATAATGAAGAGTCCTGTAGGCACGCAGTGAATATGATCTGCGACAAGTACGAGCGATCTTTGACAGGTATGGAGGT
               I  Q  L  L  A  I  V  P  D  N  E  E  S  C  R  H  A  V  N  M  I  C  D  K  Y  E  R  S  L  T  G  M  E  V

                       1110      1120      1130      1140      1150      1160      1170      1180      1190      1200
               ....|....|....|....|....|....|....|....|....|....|....|....|....|....|....|....|....|....|....|....|
RferWhite_ORF  AACATTGGAGTCTAGTACTAAGGTCGGAGACTTTCTTAAGAATACCGAGACCGAGTTGTGGCTGAACGGGACCTCTAACGGCCGCAAAAGTCCTTACAAA
                 T  L  E  S  S  T  K  V  G  D  F  L  K  N  T  E  T  E  L  W  L  N  G  T  S  N  G  R  K  S  P  Y  K

                       1210      1220      1230      1240      1250      1260      1270      1280      1290      1300
               ....|....|....|....|....|....|....|....|....|....|....|....|....|....|....|....|....|....|....|....|
RferWhite_ORF  GCGTCTTGGTGCGCTCAGTTCAGGGCCGTCTTATGGAGATCGTGGTTGAGCATCCTGAAGGAGCCACTCCTCGTTCGAGTACGCTTATTACAGACAATAT
                A  S  W  C  A  Q  F  R  A  V  L  W  R  S  W  L  S  I  L  K  E  P  L  L  V  R  V  R  L  L  Q  T  I

                       1310      1320      1330      1340      1350      1360      1370      1380      1390      1400
               ....|....|....|....|....|....|....|....|....|....|....|....|....|....|....|....|....|....|....|....|
RferWhite_ORF  TGGTATCTTTAATACTCGGATCTATTTACTACGGTCAGGTCGTAAACCAGGACGGCGTTATGAACATCAATGGCGTCCTGTTTATTTTTCTCACCAACTT
               L  V  S  L  I  L  G  S  I  Y  Y  G  Q  V  V  N  Q  D  G  V  M  N  I  N  G  V  L  F  I  F  L  T  N  L

                       1410      1420      1430      1440      1450      1460      1470      1480      1490      1500
               ....|....|....|....|....|....|....|....|....|....|....|....|....|....|....|....|....|....|....|....|
RferWhite_ORF  GACATTCCAGAACGTCTTCGCCGTAATAAACGTGTTTTCGGCTGAATTACCTCTGTTCTTGCGTGAGCACAAAAATGGCATGTACAGGACCGACGTATAT
                 T  F  Q  N  V  F  A  V  I  N  V  F  S  A  E  L  P  L  F  L  R  E  H  K  N  G  M  Y  R  T  D  V  Y

                       1510      1520      1530      1540      1550      1560      1570      1580      1590      1600
               ....|....|....|....|....|....|....|....|....|....|....|....|....|....|....|....|....|....|....|....|
RferWhite_ORF  TTTCTTGGAAAAACGATAGCGGAAATACCGTTTTTCGTTTTCCTTCCCCTAGTTTTTATATCGATATGTTATTTTCTCATCGGCCTGAATAGCGAAATGC
                F  L  G  K  T  I  A  E  I  P  F  F  V  F  L  P  L  V  F  I  S  I  C  Y  F  L  I  G  L  N  S  E  M

                       1610      1620      1630      1640      1650      1660      1670      1680      1690      1700
               ....|....|....|....|....|....|....|....|....|....|....|....|....|....|....|....|....|....|....|....|
RferWhite_ORF  CCAGATTCTTTGTGGCTTGCGGCATCGTTGTTCTGGTAGCCAATGCAGCAACAAGTTTTGGTTATTTGATATCATGCCTGTCTTCAAGTGTTTCGATGGC
               P  R  F  F  V  A  C  G  I  V  V  L  V  A  N  A  A  T  S  F  G  Y  L  I  S  C  L  S  S  S  V  S  M  A

                       1710      1720      1730      1740      1750      1760      1770      1780      1790      1800
               ....|....|....|....|....|....|....|....|....|....|....|....|....|....|....|....|....|....|....|....|
RferWhite_ORF  CCTGTCAATCGGACCACCTTTAATAATGCCCTTCTTGCTATTTGGAGGATTCTTCTTAAACATCGATTCTATACCGATCTATTTCGAATGGTTGTCGTAC
                 L  S  I  G  P  P  L  I  M  P  F  L  L  F  G  G  F  F  L  N  I  D  S  I  P  I  Y  F  E  W  L  S  Y

                       1810      1820      1830      1840      1850      1860      1870      1880      1890      1900
               ....|....|....|....|....|....|....|....|....|....|....|....|....|....|....|....|....|....|....|....|
RferWhite_ORF  TTCTCCTGGTTTAAGTACGGCAATGAGGCCCTGCTCATTAACCAATGGCAAAACATGACTGATATACAGTGTGATGAGAACAGCCGAAACTGTCCAAGAA
                F  S  W  F  K  Y  G  N  E  A  L  L  I  N  Q  W  Q  N  M  T  D  I  Q  C  D  E  N  S  R  N  C  P  R

                       1910      1920      1930      1940      1950      1960      1970      1980      1990      2000
               ....|....|....|....|....|....|....|....|....|....|....|....|....|....|....|....|....|....|....|....|
RferWhite_ORF  ATGGACACGTTGTTTTGGAAATGTATAATTTCAAAGAAGACGACTTCTTCCTGGATATATACGCCATGGTAGGTTTGATCCTGTTTTTCAGATTAGCAGC
               N  G  H  V  V  L  E  M  Y  N  F  K  E  D  D  F  F  L  D  I  Y  A  M  V  G  L  I  L  F  F  R  L  A  A

                       2010      2020      2030      2040
               ....|....|....|....|....|....|....|....|
RferWhite_ORF  GTTTTTCGTACTGTTAAGGAGAACCTATAAGGACCAATGA
                 F  F  V  L  L  R  R  T  Y  K  D  Q  *
```

**Figure S1a:** *R. ferrugineus white* (RferWhite) protein motif and domain profile.

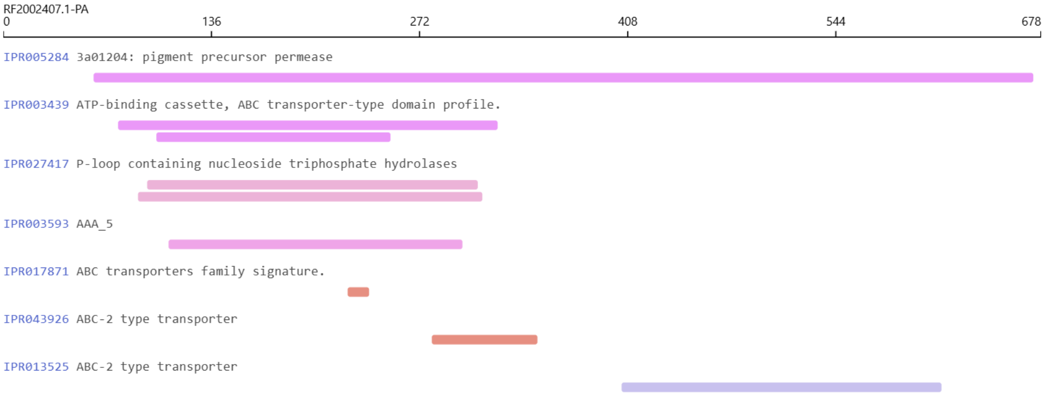


**Figure S2** RferCar Open Reading Frame (ORF). The region corresponding to the guide RNA (underlined) and the PAM site (dotted line) are shown.

```
                    10        20        30        40        50        60        70        80        90       100
             ....|....|....|....|....|....|....|....|....|....|....|....|....|....|....|....|....|....|....|....|
RferCar_ORF  ATGTCTCAAAGTCAATTTAATGAAAGAACCAGACTGTTAGGTGCACAACAGGACCAATCAATATACGAACATAATTTATTAAGAGATAGAAAAAGAAGAG
             M  S  Q  S  Q  F  N  E  R  T  R  L  L  G  A  Q  Q  D  Q  S  I  Y  E  H  N  L  L  R  D  R  K  R  R

                   110       120       130       140       150       160       170       180       190       200
             ....|....|....|....|....|....|....|....|....|....|....|....|....|....|....|....|....|....|....|....|
RferCar_ORF  TGAAACAATTCCAGTGTTGTTTATGTACAATTATTATAATTTTTGTTTTAACCATTCTTATAATAACTATATCTTACTTTAACAATTTGGCTCCGAAAGA
             V  K  Q  F  Q  C  C  L  C  T  I  I  I  I  F  V  L  T  I  L  I  I  T  I  S  Y  F  N  N  L  A  P  K  D

                   210       220       230       240       250       260       270       280       290       300
             ....|....|....|....|....|....|....|....|....|....|....|....|....|....|....|....|....|....|....|....|
RferCar_ORF  TTCAAACAATACCCAATCCAACAATGAATCTTCATCAAATAATTTACCCGCTGACACATTTATAAAATTGCATCAAGTATTATCATGGCCTAGAACAGAT
              S  N  N  T  Q  S  N  N  E  S  S  S  N  N  L  P  A  D  T  F  I  K  L  H  Q  V  L  S  W  P  R  T  D

                   310       320       330       340       350       360       370       380       390       400
             ....|....|....|....|....|....|....|....|....|....|....|....|....|....|....|....|....|....|....|....|
RferCar_ORF  AAAATTGACCCAGTAAGTTATAAAGATTATAATGAGTTAAATAAAGCTATAACAGAAGGTAGAAAAGCCTTAAATAGCAAATATGAATCTGAAAAATACA
              K  I  D  P  V  S  Y  K  D  Y  N  E  L  N  K  A  I  T  E  G  R  K  A  L  N  S  K  Y  E  S  E  K  Y

                   410       420       430       440       450       460       470       480       490       500
             ....|....|....|....|....|....|....|....|....|....|....|....|....|....|....|....|....|....|....|....|
RferCar_ORF  ACAGCCCATTAAAGTTAAATAGTCCAAGTTATAAACATCAAAAAGTGATGGCTACTAGTGAAAAGGCCAGAAGTATATCACAAAAAGCTTTTGAAGGAGA
             N  S  P  L  K  L  N  S  P  S  Y  K  H  Q  K  V  M  A  T  S  E  K  A  R  S  I  S  Q  K  A  F  E  G  E

                   510       520       530 gRNA2 540       550       560       570       580       590       600
             ....|....|....|....|....|....|....|....|....|....|....|....|....|....|....|....|....|....|....|....|
RferCar_ORF  AGGAGCTACAAAGTATTTTTACAGAAATTGGAACAAATACACACCAGTAGGTAAAATTTGCTTTACCAACATTACAGATTCAGTACTAACGGACTTATGC
              G  A  T  K  Y  F  Y  R  N  W  N  K  Y  T  P  V  G  K  I  C  F  T  N  I  T  D  S  V  L  T  D  L  C

                   610       620       630       640       650       660       670       680       690       700
             ....|....|....|....|....|....|....|....|....|....|....|....|....|....|....|....|....|....|....|....|
RferCar_ORF  TATGATGCAATACCGTGCAATTTTTCTTATAAATACAGATCTTACAATGGTTCATGTAATAATGTTAATCACCCTAACAGCTACGGAGTTGCTTATAGGC
              Y  D  A  I  P  C  N  F  S  Y  K  Y  R  S  Y  N  G  S  C  N  N  V  N  H  P  N  S  Y  G  V  A  Y  R

                   710       720       730       740       750       760       770       780       790       800
             ....|....|....|....|....|....|....|....|....|....|....|....|....|....|....|....|....|....|....|....|
RferCar_ORF  CATTTAGGCGAGCTTTACCACCAGATTATGCTGACGGTATAAGTAAACCTAGAGTGTCTACATTTGGCGATAATCTACCGTCTGCCCGTAGAGTCAGTGT
             P  F  R  R  A  L  P  P  D  Y  A  D  G  I  S  K  P  R  V  S  T  F  G  D  N  L  P  S  A  R  R  V  S  V

                   810       820       830       840       850       860       870       880       890       900
             ....|....|....|....|....|....|....|....|....|....|....|....|....|....|....|....|....|....|....|....|
RferCar_ORF  AGAAATACATCGACCTTATTACAGAGACGACTCCAAGTTTTCAGTTATGTTAGCGGTTTGGGGTCAATTTCTCGACCATGACGTTACTGCTACAGCACTT
              E  I  H  R  P  Y  Y  R  D  D  S  K  F  S  V  M  L  A  V  W  G  Q  F  L  D  H  D  V  T  A  T  A  L

                   910       920       930       940       950       960       970       980       990      1000
             ....|....|....|....|....|....|....|....|....|....|....|....|....|....|....|....|....|....|....|....|
RferCar_ORF  AGTCAAAAACAGAATGGTGAAAGCATATCCTGCTGTAACTCTAATGATACCTTACCTTCCGAATGTTTTCCTGTTTACTTGGATATCAATGATTATTTTC
              S  Q  K  Q  N  G  E  S  I  S  C  C  N  S  N  D  T  L  P  S  E  C  F  P  V  Y  L  D  I  N  D  Y  F

                  1010      1020      1030      1040      1050      1060      1070      1080      1090      1100
             ....|....|....|....|....|....|....|....|....|....|....|....|....|....|....|....|....|....|....|....|
RferCar_ORF  AAAAATTTAATATATCCTGCATGGAGTTTGTTAGATCTGCCCCAGCACCAACATGTTGTTTAGGACCAAGAGAACAGATGAATCAAGTTACTGCTTTCAT
             Q  K  F  N  I  S  C  M  E  F  V  R  S  A  P  A  P  T  C  C  L  G  P  R  E  Q  M  N  Q  V  T  A  F  I

                  1110      1120      1130      1140      1150      1160      1170      1180      1190      1200
             ....|....|....|....|....|....|....|....|....|....|....|....|....|....|....|....|....|....|....|....|
RferCar_ORF  AGATGGTTCTGTAGTATATGGAGCTGATGAAAATACAGTAAAATCATTAAGAACCATGGAAAAAGGTTTGCTGAAAATGTTTATTACTAGTGATAACAGA
              D  G  S  V  V  Y  G  A  D  E  N  T  V  K  S  L  R  T  M  E  K  G  L  L  K  M  F  I  T  S  D  N  R

                  1210      1220      1230 gRNA1 1240      1250      1260      1270      1280      1290      1300
             ....|....|....|....|....|....|....|....|....|....|....|....|....|....|....|....|....|....|....|....|
RferCar_ORF  TCATTACTTCCAGCATCTTATGATATGACTGATGGTTGTAACCGGGAGGAGCAAAAAAATAAAGGAAGATATTGCTTTTTGTCTGGTGATAAACGAGCAA
              S  L  L  P  A  S  Y  D  M  T  D  G  C  N  R  E  E  Q  K  N  K  G  R  Y  C  F  L  S  G  D  K  R  A

                  1310      1320      1330      1340      1350      1360      1370      1380      1390      1400
             ....|....|....|....|....|....|....|....|....|....|....|....|....|....|....|....|....|....|....|....|
RferCar_ORF  ATGAAAATCTTCATTTAACAAGCATGCACTTGATATGGGCAAGACAACACAATTTAATAGCAAAAAATCTTGCTCAATTAAATCCCCACTGGAATGATGA
             N  E  N  L  H  L  T  S  M  H  L  I  W  A  R  Q  H  N  L  I  A  K  N  L  A  Q  L  N  P  H  W  N  D  E

                  1410      1420      1430      1440      1450      1460      1470      1480      1490      1500
             ....|....|....|....|....|....|....|....|....|....|....|....|....|....|....|....|....|....|....|....|
RferCar_ORF  AACAATTTTTCAAGAATCTAGACGTATTTTGTCAGCTCAAATGCAACATATTACGTATAATGAATTTCTACCGATTCTATTGGGTAAACATGTGATGGAA
              T  I  F  Q  E  S  R  R  I  L  S  A  Q  M  Q  H  I  T  Y  N  E  F  L  P  I  L  L  G  K  H  V  M  E

                  1510      1520      1530      1540      1550      1560      1570      1580      1590      1600
             ....|....|....|....|....|....|....|....|....|....|....|....|....|....|....|....|....|....|....|....|
RferCar_ORF  AAATATGAGCTATCACCGAAGCAGCAAGGTTATTTTTACAAATATAACCCAACTATTGACCCTAGCGTTGCAAATGTTTTCGCAGCATCAGCTTTCCGTT
              K  Y  E  L  S  P  K  Q  Q  G  Y  F  Y  K  Y  N  P  T  I  D  P  S  V  A  N  V  F  A  A  S  A  F  R

                  1610      1620      1630      1640      1650      1660      1670      1680      1690      1700
             ....|....|....|....|....|....|....|....|....|....|....|....|....|....|....|....|....|....|....|....|
RferCar_ORF  TTGCCCATTCATTAATACCAACTCTAATTAAAATACTAGCTAATCACTCGAATAGTACTGAATATATCCAGATGCATAAAATGCTTTTCGATCCTTTTAA
             F  A  H  S  L  I  P  T  L  I  K  I  L  A  N  H  S  N  S  T  E  Y  I  Q  M  H  K  M  L  F  D  P  F  K

                  1710      1720      1730      1740      1750      1760      1770      1780      1790      1800
             ....|....|....|....|....|....|....|....|....|....|....|....|....|....|....|....|....|....|....|....|
RferCar_ORF  ACTATACGATAATGGCGAATTAAATAAAGCTTTAAACGGGGCAATGAATACTCATGTTGAAGCAAGCGATGCTTATTTTTCCAAGGAAATAAAATTTCAT
              L  Y  D  N  G  E  L  N  K  A  L  N  G  A  M  N  T  H  V  E  A  S  D  A  Y  F  S  K  E  I  K  F  H
```

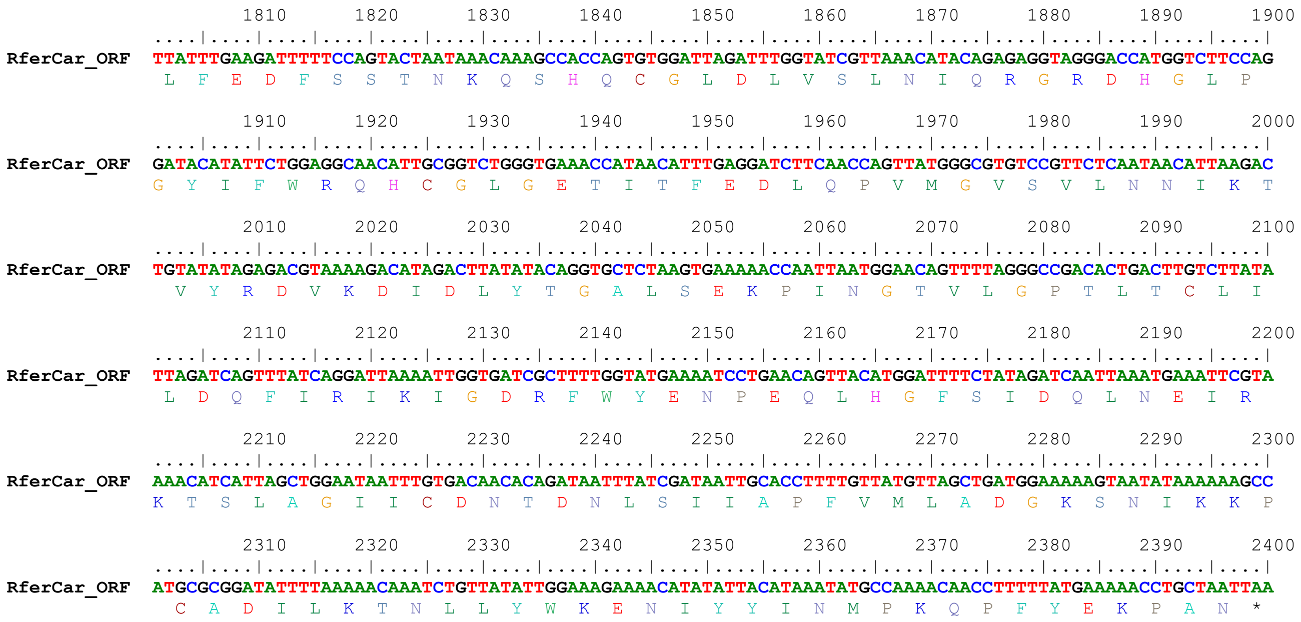


**Figure S2a:** *R. ferrugineus cardinal* (RferCar) protein motif and domain profile.

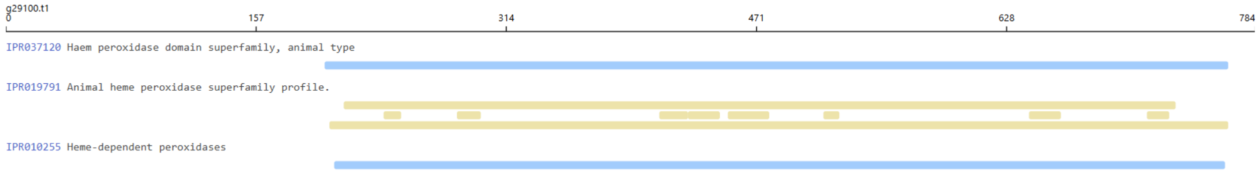


**Figure S3**. RferWhite and RferCar tissue specific expression gel images. The tissues in wells are represented in the order as M (marker), A (antennae), G (gut and fatdody), H (head), L (leg), T (thorax), and W (wings) (see Figure 2D for details).

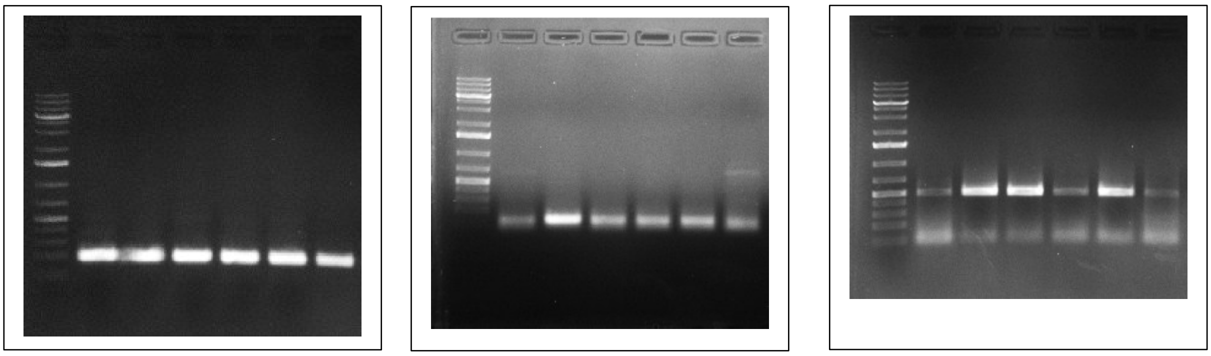

**Figure S4**. Primer walking (using RfCar-GT-R1) (Table S2) Sanger sequence chromatogram (anti-sense strand) of RferCar ORF region corresponding to the guide RNA of *cardinal* mutant 13-nt homozygous RPWs. The arrow indicates the deleted 'GTTACAACCATCA' sequence from the gRNA region and the PAM shown in the box.

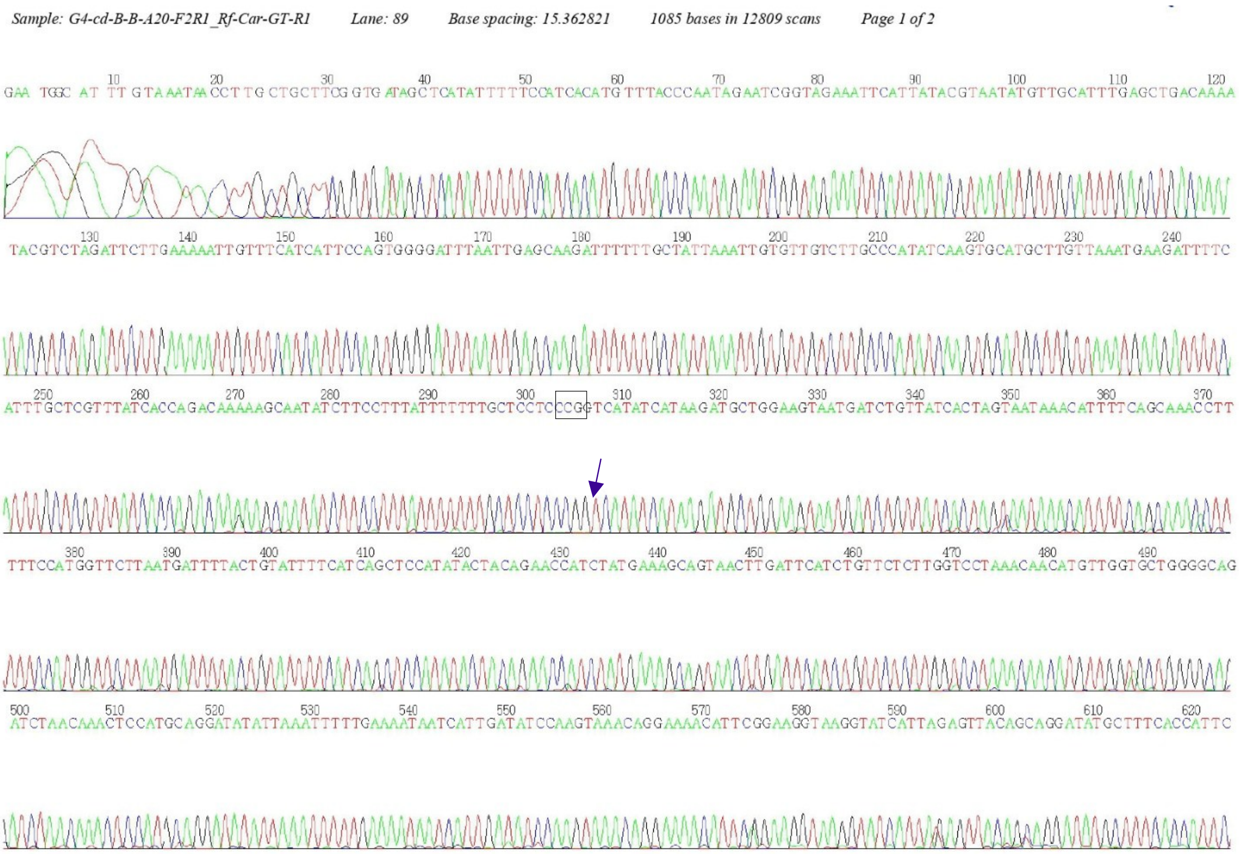
